# HSA^+^ immature cardiomyocytes persist in the adult heart and expand after ischemic injury

**DOI:** 10.1101/518530

**Authors:** Mariana Valente, Tatiana Pinho Resende, Diana Santos Nascimento, Odile Burlen-Defranoux, Benoit Dupont, Ana Cumano, Perpétua Pinto-do-Ó

**Affiliations:** i3S – Instituto de Investigação e Inovação em Saúde, Universidade do Porto, 4200-135, Porto, Portugal.; INEB – Instituto Nacional de Engenharia Biomédica, Universidade do Porto, 4200-135,Porto, Portugal.; ICBAS – Instituto de Ciências Biomédicas Abel Salazar, Universidade do Porto, 4050-313, Porto, Portugal.; Unit for Lymphopoiesis, Immunology Department, INSERM U1223, Institut Pasteur, 75015, Paris, France.; Université Paris Diderot, Sorbonne Paris Cité, Cellule Pasteur, 75018, Paris, France.; Beckman Coulter, France S.A.S, 93420 Villepinte, France.

## Abstract

The assessment of the regenerative capacity of the heart has been compromised by the lack of surface signatures to characterize cardiomyocytes. Here, combined multiparametric surface marker analysis with single cell transcriptional profiling and in vivo transplantation, identify the main fetal cardiac populations and their progenitors. We found that cardiomyocytes at different stages of differentiation co-exist during development. We identified a population of immature HSA/CD24^+^ cardiomyocytes that persists throughout life and that, unlike other cardiomyocyte subsets, actively proliferates up to one week of age and engraft cardiac tissue upon transplantation. In adult heart HSA/CD24^+^ cardiomyocytes appear as mononucleated cells that cycle and increase in frequency after infarction. Our work identified cell surface signatures that allow the prospective isolation of cardiomyocytes at any developmental stage and the detection of adult cardiomyocytes poised for activation in response to ischemic stimuli. This work opens new perspectives in the understanding and treatment of heart pathologies.

## Introduction

The cell types that form the mammalian heart have diverse developmental origins and temporal differentiation^1^. In the mouse, cardiomyocytes (CMs) are initially specified by embryonic day (E) 7.5 from the first set of cardiogenic progenitors (first heart field)^2^ followed by the incoming cells from the second heart field (SHF)^3^. At E 9.5 (looping-heart stage), the heart is divided into primitive (*P*) atria (*At*), ventricle *Vt (PVt*) and outflow track (*OFT*, future great vessels and atrioventricular junction *GV-AVJ*) and is composed of CMs and endocardial cells (EndoCs) that form the endocardium. Cells migrating from the peripheral tissues shape the final heart morphology. From E9.5 to E11.0, epicardial cells (EpiCs) derive from the proepicardial organ and coat the heart surface (epicardium). A fraction of EpiCs undergo epithelial-to-mesenchymal transition (EMT), giving rise to epicardial-derived cells (EPDCs), which migrate into the myocardium and generate peri-vascular smooth muscle cells (SMCs)^4^ and interstitial fibroblasts (FBs)^5,6^.

CMs contribute to heart growth^7,8^ through extensive cell divisions, exiting cell cycle as development progresses, a process virtually completed by the end of the first week of postnatal life^9^. The CM compartment enlarges thereafter by hypertrophy that in rodents coincides with binucleation of myocytes^9,10^.

Several markers have been individually used to identify CMs (SIRPα, VCAM-1, Caveolin-3 (Cav3)^11^-^13^) and their putative adult progenitors (Sca-1^14,15^, c-kit^16^), FBs (Ddr2, Thy1^17,18^), SMCs (PDGFrβ^19^) and ECs (PECAM-1^20^). However, because each of these markers is expressed in other cell types, including circulating and heart resident hematopoietic cells, they do not, when used alone, unambiguously define cardiac progenitors (PRGs) or mature populations.

Myocardium regeneration requires the production of new adult CMs. Consistent with a potential regenerative capacity of the adult heart, CM replacement by expansion, albeit at low rate, has been recently reported indicating that not all adult CMs are post-mitotic cells^21,22^. However, the efforts to unravel mechanism/s of neo-cardiomyogenesis have been hampered by the lack of strategies to identify and prospectively isolate the rare CM capable of turnover in normal and diseased hearts.

To identify different maturation stages of CMs and to follow their development up to adulthood we analyzed the phenotype of all cells in the developing mouse heart. Multi-parametric flow cytometry combined with single cell multiplex qRT-PCR of purified cell subsets allows the identification of distinct cell types and the definition of their lineage affiliation. HSA (CD24) expression is consistently associated with the CM lineage and, combined with other surface proteins, unraveled coexisting subsets of CMs at different stages of maturation. The most immature population expresses HSA, ALCAM, MCAM and troponin T (Tnnt), but not Cav3 and is capable to integrate heart tissue, after transplantation. The progressive loss of those surface markers contemporaneous with the expression of Cav3 and binucleation identifies mature CMs. Importantly, HSA^+^, but not HSA^-^ Cav3^+^ CM, isolated throughout development and up to P7 actively proliferate and spontaneously acquire contractile properties *in vitro*. Isolated from E15, they engraft heart tissue transplanted in the ear pinna of adult mice and persist, albeit at low frequencies, in adult hearts. This population responds to myocardial infarction (MI) by proliferating and increasing in numbers.

## Results

### Phenotype assignment to the cellular components in the fetal heart

To resolve the phenotype of the cellular components of the developing heart, we screened by flow cytometry cell suspensions isolated from the three heart regions (i.e. *At, GV-AVJ* – containing the connection of the four cavities, the great vessels and the valves – and *Vt*, Fig. 1a) for the expression and relative abundance of 30 surface proteins. The analysis was performed in E17.5 hearts that have similar structure and cellular components to those observed in the adult. We selected 11 antibodies recognizing surface proteins from which HSA (CD24) had not been previously associated with cardiac cells. Following a sequential gating strategy (Fig. 1a; Supplementary Fig. 1), we identified 13 distinct cell populations, after exclusion of hematopoietic (CD45^+^Ter119^+^) cells. Multiparametric analysis of the flow cytometry data by non-linear dimensionality reduction algorithms (Fig. 1b) created maps where cells clustered together according to phenotype (t-SNE, upper graphs) or were organized into hierarchies of related phenotypes (spanning tree, lower graphs). ICAM-1^+^ cells (olive arrow) were predominant in the *At*, Sca-1^+^ (blue arrow) and ALCAM^+^ (cyan asterisk) in the *GV-AVJ* and Thy1^+^ (green arrow) in the *Vt*. The remaining subsets were represented in all three regions, albeit at different frequencies (Fig. 1b).

**Fig. 1.**
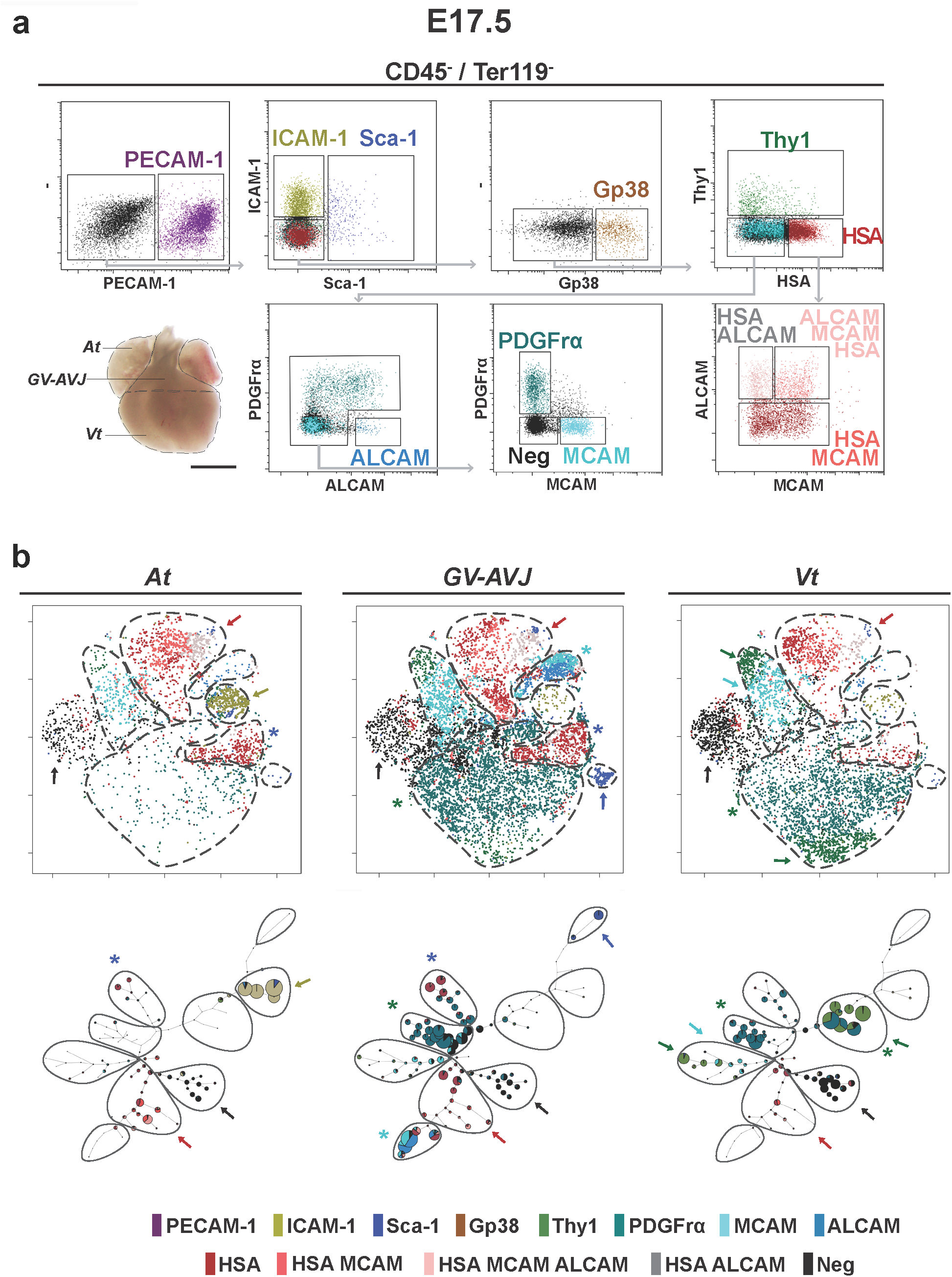
Cell subsets in fetal heart. **a**. Dissected cardiac regions in E17.5 fetal heart: *At, GV-AVJ* and *Vt*. Scale bar: 1 mm. Flow cytometry dot plots of *At* CD45^-^Ter119^-^cells showing the expression of PECAM-1, ICAM-1, Sca-1, Gp38, Thy1, HSA, PDGFrα, ALCAM and MCAM (n=10). **b.** t-SNE (upper graphs) and Spanning Tree (lower graphs) analyses applied to the populations defined in A. Each point represents a cell and colors represent surface signatures (see Supplementary Fig. 1). Marker expression is represented by a color code: red arrows HSA, MCAM and ALCAM, cyan arrows MCAM, green asterisks PDGFrα, olive arrows ICAM-1, green arrows Thy1, blue asterisks HSA, blue arrow Sca-1, cyan asterisk ALCAM and black arrows none of the above. Colors correspond to populations as in **a**. and number of dots denotes relative size of the populations. Multiple colors in the same node represent co-expression. *B*. *At* | Atria; *GV-AVJ* | Great Vessels-Atrioventricular Junction; *Vt* | Ventricles.

To determine the cell identity of the newly defined populations, we analyzed their transcriptional profiles and anatomical distribution. The expression levels of 31 transcripts affiliated to different cardiac lineages were analyzed in purified cells (20 cells) from each subset. Unsupervised hierarchical clustering and principal component analysis (PCA) grouped the subsets in seven clusters (Fig. 2a-b; Supplementary Table 1). We observed a strong correlation between the clustering by the transcriptional profiles and the subset definition using the cell surface markers, highlighting the validity and the robustness of our approach. Cluster I encompassed CMs identified by the expression of *Nkx2-5, Tnnt2* and *Des*. *At* CMs (Cluster I*a*) expressed *At* specific myosin, *Myl7* and *Myh6* and were phenotypically characterized by the co-expression of HSA, MCAM and ALCAM. *Vt* CMs (Cluster I*b*) expressing the *Vt* myosin *Myl2* and *Myh7*, expressed HSA and MCAM, but not ALCAM (Fig. 2a-b). We also analyzed the expression and distribution of the identified proteins *in situ (*Fig. 3a). HSA^+^CMs identified by the characteristic striated Actinin pattern were found in both chambers (more frequent in the *At* than in the *Vt*, Fig. 3f). In cluster II cells expressed *Acta2*, indicative of their SMC affiliation, together with *Myh11* and corresponded to ALCAM^+^ *GV-AVJ* cells found in the wall of the great vessels, co-expressing SMA protein (Fig. 2a-b; 3d). Cluster III cells expressed *Kdr, Flk1* and *Tek* and comprised PECAM-1^+^ ECs and EndoCs (Fig. 2a-b), delineating the blood vessels (Fig. 3c-e) and the inner surface of the myocardium (EndoC Cluster IV cells expressed *Wt1, Tbx18, S100a4* and *Gja1*, were identified as ICAM-1^+^ (Fig. 2a-b) and were observed in the sub-epicardial region of both chambers (Fig. 3b, e (inset #)), supporting their EPDCs identity. Cluster V combined cells exhibiting a FB transcriptional profile (*Col1a1, Col3a1, Dcn, Twist1, Ddr2, Vcan* and *Fn1*) with the expression of *GV-AVJ* specific genes *Tbx3* together with *Isl1*, revealing a Sca-1 or HSA phenotype (Fig. 2a-b). Sca-1^+^ (PECAM-1^-^) cells were detected at the insertion of great vessels (Fig. 3c, *), whereas HSA^+^ cells (Actinin^-^) were observed in the EndoC cushion mesenchyme (Fig. 3c, *). Cluster VI corresponded to MCAM^+^ cells expressing the SMC-associated transcript *Acta2* together with *Tbx20, Snai2, Vim, Fn, Fap (*Fig. 2a-b). Finally, in Cluster VII, Gp38^+^, PDGFrα^+^ and Thy1^+^ cells were grouped by the expression of a different combination of FB-related transcripts (*Ddr2, Col3a1, Dcn, Postn, Fn, Vim, Snai2, Tbx20*) (Fig. 2a-b). PDGFrα expression was associated with interstitial FBs in the myocardium (Fig. 3b), while Gp38^+^ cells were detected in the epicardium (Fig. 3e, *).

**Fig. 2.**
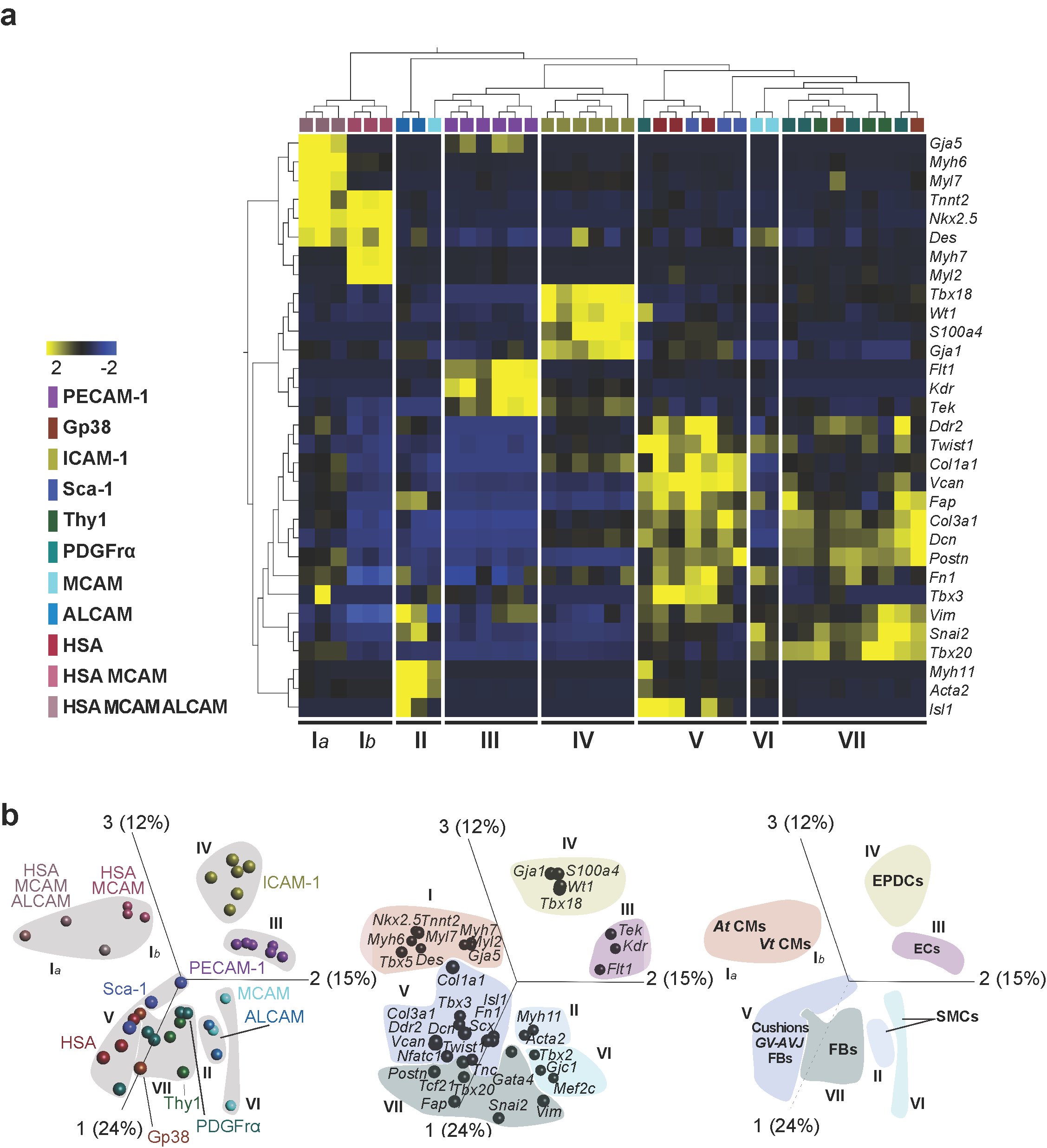
Transcriptional profiles assign cardiac lineages to phenotypes. **a.** Unsupervised hierarchical clustering of multiplex qRT-PCR data (20 sorted cells, n=3) in E17.5 hearts (acronyms and color-code as in Supplementary Fig. 1). Differential expressed genes among clusters assign a cell type to each surface signature. Statistical significance was determine using two-way ANOVA test; *p*=0.007, *q*=0.01 (Table 1). **b.** Principal component analysis (PCA) of the transcriptional profile in **a**. clustered by surface phenotype (left graph), by gene expression (middle graph) or by cardiac cell type defined by gene expression (right graph). Cluster I*a* – *At* CMs, Cluster I*b* – *Vt* CMs, Cluster II and VI – SMCs, Cluster III – ECs, Cluster IV – EPDCs, Cluster V – Cushions *GV-AVJ* FBs and Cluster VII FBs. *At* CMs | Atria Cardiomyocytes; *Vt* CMs | Ventricles Cardiomyocytes; ECs | Endothelial Cells; EPDCs | Epicardial-Derived Cells; SMCs | Smooth Muscle Cells; FBs | interstitial Fibroblasts; *GV-AVJ* FBs | Great Vessels-Atrioventricular Junction Fibroblasts.

**Fig. 3.**
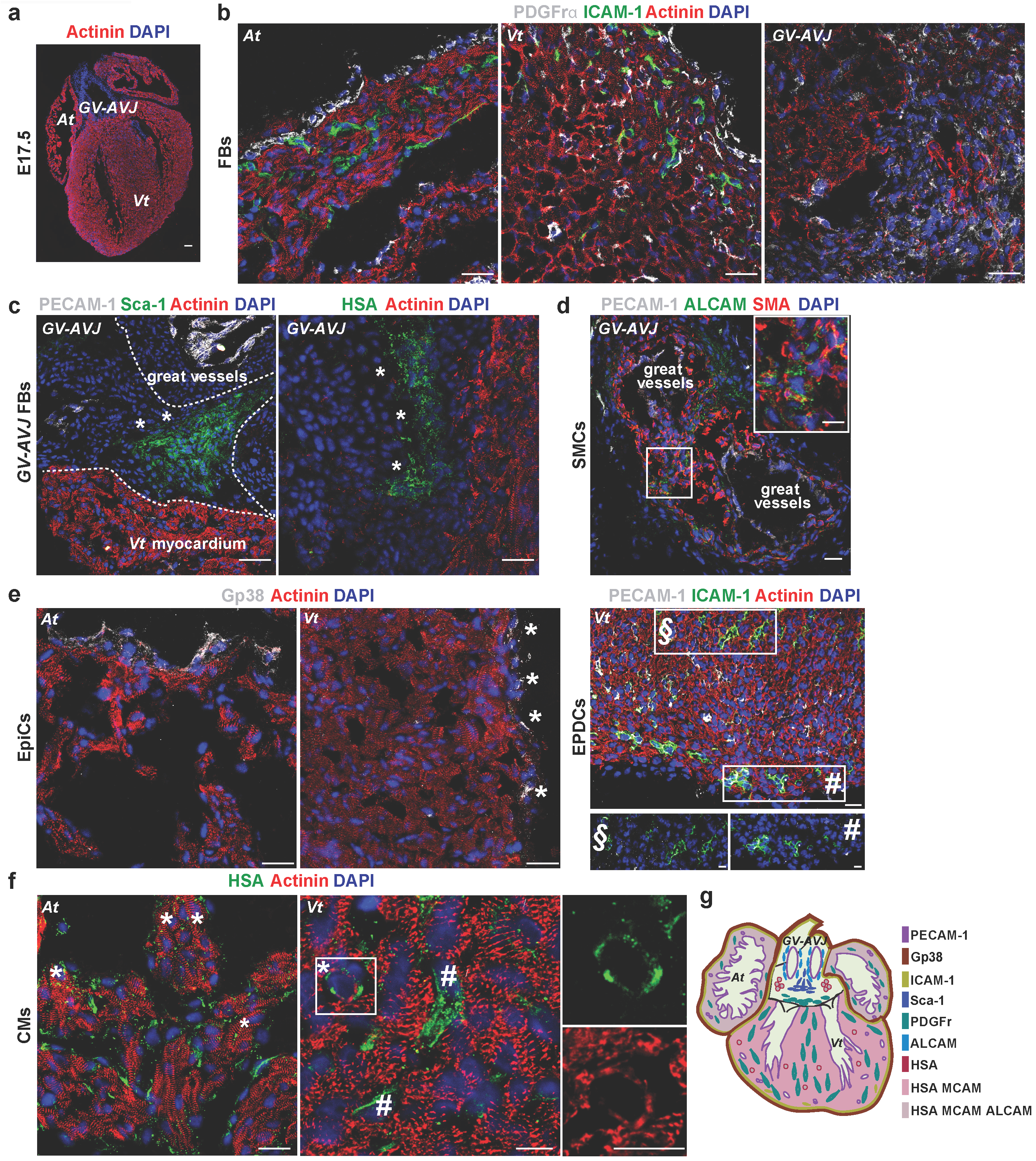
Spatial distribution of cardiac populations. **a.** Coronal view of E17.5 heart section: Actinin (red), nuclear content (DAPI, blue). Scale bar: 50 µm. **b.** Interstitial FBs (PDGFrα^+^ cells). **c.** *GV-AVJ* FBs (*, Sca-1^+^PECAM-1^-^ or HSA^+^Actinin^-^ cells); **d.** SMCs (ALCAM^+^SMA^+^ cells, inset); **e.** EpiCs (two left panels, Gp38^+^ cells, *); EPDCs (right panel, ICAM-1^+^ PECAM-1^-^, Inset #.; ECs, ICAM-1^+^PECAM-1^+^ cells in inset §.; **f.** CMs (HSA^+^Actinin^+^, *; stromal cells, HSA^+^Actinin^-^ cells in the *Vt* #). Scale bar: 20 µm, insets: 10 µm. **g.** Fetal heart representation with the spatial distribution of the newly defined cardiac populations. Color-code as in Supplementary Fig. 1. CMs | Cardiomyocytes; EpiCs | Epicardial Cells; EPDCs | Epicardial-Derived Cells; ECs | Endothelial Cells; SMCs | Smooth Muscle Cells; FBs | interstitial Fibroblasts; *GV-AVJ* FBs | Great Vessels-Atrioventricular Junction Fibroblasts; *At* | Atria; *Vt* | Ventricles; *AVJ* | Atrioventricular Junction.

To probe the homogeneity of the cardiac subsets here defined we carried out single-cell transcriptional analyses (Supplementary Fig. 2a-b). Cell sorting was performed using the index-sorting tool that records, for each cell, the levels of expression of each phenotypic parameter (Supplementary Fig. 2c). Both the heat-map and the PCA analysis indicated that the phenotypically defined populations homogeneously co-expressed the analyzed transcripts, underscoring the strength of this combined approach (Supplementary Fig. 2a-b).

The ensemble of these results validated a phenotyping strategy (Fig. 1; Supplementary Table 1) that allowed the identification (Fig. 2a-b) and the prospective isolation of the major cardiac cell subsets in the fetal heart (Fig. 3g).

### Different stages of CM maturation co-exist during development

A CM transcriptional profile (*Nkx2-5, Tnnt2* and *Des*) was associated with membrane HSA expression in different, but related, cell populations (Fig. 2) isolated from both heart chambers. To understand the kinetics of the newly defined populations along heart morphogenesis we performed a similar analysis at earlier developmental stages (E9.5 and E13.5, Supplementary Fig. 33, c, d). HSA^+^MCAM^+^ALCAM^+^ (triple positive) CMs were detected in all analyzed embryonic stages. However, by E17.5 most *Vt*, but not *At*, HSA^+^ CMs had lost ALCAM expression (Supplementary Fig. 3a). HSA^+^MCAM^+^(ALCAM^-^) and HSA^+^(MCAM^-^ALCAM^-^) CMs were more frequent in the E17.5 *Vt (*Supplementary Fig. 33, bottom right plot) and triple negative CMs (HSA^-^ MCAM^-^ALCAM^-^), initially detected in the E13.5 *Vt*, increased to comprise most CMs at E17.5 (Fig. 4a, d; Supplementary Fig. 33, c, d). Cell cycle analysis showed that E9.5 HSA^+^ cells, with a transcriptional profile of CM PRGs (Fig.4b), were highly proliferative (63.7% in G1 and 15.9% in S/G2-M). As development proceeded, the frequency of HSA^+^ cells decreased, and they also divided less (14.4% versus 7.66% were in S/G2-M at E13.5 and E17.5, respectively, Supplementary Fig. 3b).

**Fig. 4.**
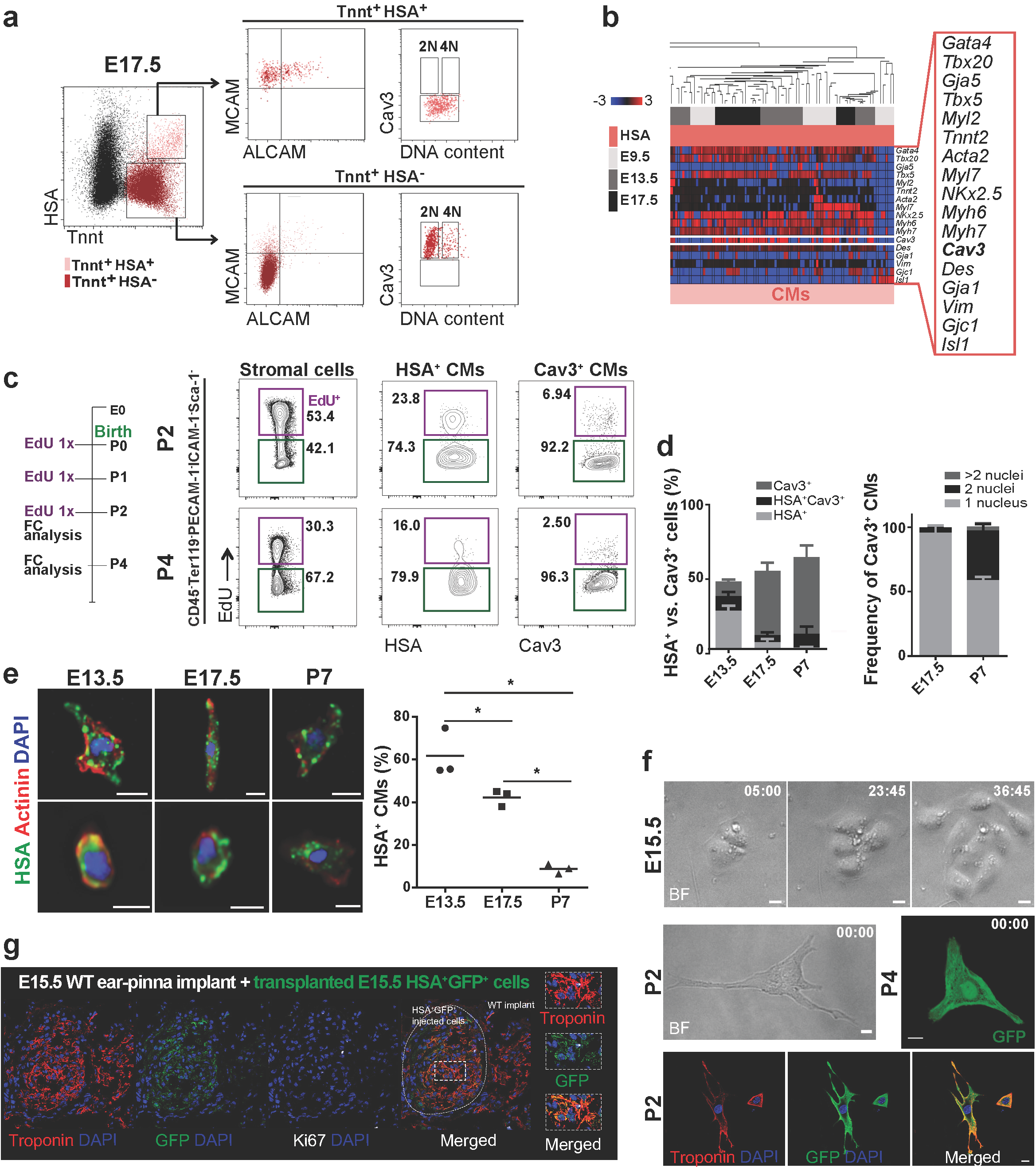
Expression of HSA and Cav3 identify two cardiomyocyte subsets. **a.** E17.5 whole heart suspensions (n=2). HSA^+^MCAM^+^ALCAM^+/-^Cav3^-^ (light pink, upper panels) and HSA^-^MCAM^-^ALCAM^-^ Cav3^+^ (red, lower panels) analyzed for DNA content. **b.** Unsupervised hierarchical clustering the HSA^+^CMs (single cell multiplex qRT-PCR data, at different E stages). **c.** EdU incorporation in CD45^-^ TER119^-^PECAM-1^-^ICAM-1^-^Sca-1^-^, PDGFRα^+^ stromal cells (left), HSA^+^ CMs (middle) and Cav3^+^ CMs (right) in P2 (top) and P4 (bottom) hearts (n=2). **d.** HSA and Cav3 expression in E13.5, E17.5 and P7 cardiac cells. Flow cytometry (left panels, n=2) and cytospin (right panels, n=3, 300 cells analyzed in each). Histograms correspond to cytology data. Scale bar: 20 µm. **e.** HSA expression in CMs (Actinin^+^) by imaging flow cytometry. Scale bar: 10 µm. Frequency of HSA^+^ CMs at E13.5, E17.5 and P7 (n=3, 59397, 67306 and 7088 total cells analyzed for E13.5, E17.5 and P7, respectively). **f.** Cultured HSA^+^ E15.5 CMs (time lapse, hr: min, upper panels, MS1, MS2), HSA^+^ P2 and P4 CMs (middle panels and MS3, MS4). Scale bar: 10 µm (n=3). Immunofluorescence of cultured (72 hours) P2 CMs. Troponin T (in red) and GFP (Ubiquitin-GFP cells, lower panels). **g.** Immunofluorescence of cardiac implants transplanted with Ub-GFP^+^ E15.5 HSA^+^ CMs (n=3). Cardiac troponin T (red), GFP (donor cells), Ki67 (white) and DAPI (nuclei, blue). Scale bar: 50 µm. Insets show higher magnification of the region within the dashed white rectangle (right panels). * p<0.05. CMs | Cardiomyocytes; *At* | atria; *Vt* | Ventricles.

To investigate whether the phenotypically distinct CM subsets corresponded to different stages of maturation, we included Tnnt^23^ and Cav3^12^ in our analysis. Surface Cav3 expression is detected in the adult, but undetectable in most immature CMs^12^, while Tnnt is widely expressed in the CM lineage^23,24^. The transcriptional profile of the HSA^+^ CMs along time (E9.5, E13.5 and E17.5) revealed that Cav3 starts to be transcribed at E9.5 (Fig. 4b), suggesting a lineage relationship between the two CM subsets. These results highlight the progressive loss of the surface markers ALCAM, MCAM and HSA coinciding with the transcriptional expression of Cav3 at E13.5, while at E17.5 the majority of CMs (Tnnt^+^) were HSA^-^MCAM^-^ALCAM^-^Cav3^+^ (Fig. 4a-b). During maturation, CMs lose proliferative capacity and suffer morphological alterations leading to increased sarcomere complexity, bigger cell size and binucleation^9^. Cell cycle analysis of E17.5 CM subsets (HSA^+^ and Cav3^+^) showed that HSA^+^ cells were still proliferative (*At*: 24.4% and 20.5%; and *Vt*: 35.5% and 25.3%, in G_1_ and S/G_2_-M, respectively) whereas Cav3^+^CMs were largely out of cycle (*At* 69.4% and *Vt* 78% in G_0_, Supplementary Fig. 3c). The stromal subsets were highly proliferative (*At*: 36.6% and 11.2%; and *Vt*: 27.6% and 11.3%, respectively in G_1_ and S/G_2_-M, Supplementary Fig. 3c). Binucleated cells (4N DNA) were observed in the Tnnt^+^HSA^-^Cav3^+^CM population, while Tnnt^+^HSA^+^Cav3^-^ CMs exhibited a continuum increase in DNA content, consistent with nucleic acid synthesis (Fig. 4a; Supplementary Fig. 4a-b). For the first time, binucleated CMs were found during development and can be isolated using HSA^-^Cav3^+^ as surface signature.

Our results demonstrate the coexistence of four stages of CM maturation along development (E9.5 – E17.5). Immature HSA^+^MCAM^+^ALCAM^+^ CMs are the dominant subset in the *At*. The expression of these markers is progressively lost such that at E17.5 only a small fraction of *Vt* CMs is still HSA^+^MCAM^+^ALCAM^-^, while the majority of CMs are negative for all markers, display surface Cav3 and have initiated binucleation (Fig. 4a; Supplementary Fig. 33, c). *At* CMs retained the expression of the three markers until later embryonic stages than their *Vt* counterparts (Supplementary Fig. 3a), consistent with the well-documented differences in CM maturation of the two chambers^25^.

Because HSA is the last surface protein to be lost before the acquisition of Cav3, we used these two markers to discriminate immature and mature CMs, respectively. Interestingly, HSA and Cav3 expression define two CMs subsets with different proliferative capacity, even after birth. Three EdU (P0, P1 and P2) injections in the neonates labeled 23% of HSA^+^ CMs detected at P2 and 16% at P4, indicating that this CM subset maintains proliferative activity after birth (Fig. 4c). By contrast, only 7% of Cav3^+^ CMs incorporated EdU, demonstrating a lower contribution of the mature CM subset to postnatal heart growth (Fig. 4c). Other non-CM (stromal) cells showed more than 50% EdU incorporation, compatible with their high proliferative activity at this stage (Fig. 4c). The reduced frequency of Cav3^+^ CMs at P7 detected by flow cytometry (Supplementary Fig. 3d, contour plots) reflects the sensitivity of postnatal CMs to enzymatic digestion. To overcome this, we performed a similar analysis in cells fixed before dissociation. Kinetics of HSA and Cav3 expression showed that the decrease of the former parallels an increase of the latter (Fig. 4d). At E13.5 we detected HSA+ (27.3% ± 3.3%), HSA^+^Cav3^+^ (10.3% ± 1.2%) and a few larger cells expressing Cav3 (10.3% ± 2.6%). At E17.5, the majority of the cells expressed Cav3 (44.8% ± 6.3%), a fraction of them were binucleated (4.1% ± 0.9%) and coexisted with smaller cells that were either HSA^+^ (4.9% ±1.5%) or HSA^+^Cav3^+^ (3.9% ±1.9%). After birth (P7), the majority of cells expressed Cav3 (53.1% ± 5.1%) and presented two (2 nuclei, 38.7% ± 2.5%, *, Fig. 4d) or more (>2 nuclei, 2.5% ± 0%, Fig. 4d) nuclei. CMs fixed prior to isolation were also analyzed by imaging flow cytometry, evidencing decreased percentage of immature HSA^+^ CMs during development (57.8% ± 11.3% at E13.5 and 38.3% ± 3.8% at E17.5) with a pronounced decline after birth (4.8% ± 2.3% at P7, Fig. 4e). Importantly, HSA expression was only observed in mononucleated CMs, further associating its expression with an immature phenotype (Fig. 4e).

To confirm that HSA discriminates immature renewing CM, purified HSA^+^ cells isolated from E15.5, P2 or P4 and adult (< 4 weeks old) hearts were cultured for up to one week. E15.5 HSA^+^ cells either divided (∼20%) or were contractile (∼75%) in culture, whereas no proliferation was observed in P2 or P4 cardiac cells, probably due to the rapid differentiation in culture, (Fig. 4f; Supplementary Fig. 5a; Supplementary Movie 1). Seeded P2 and P4 cells adhered and were contractile at a frequency ranging from 1:30-1:40 (Supplementary Fig. 5b), by contrast Cav3^+^ cells, irrespective of the stage at which they were isolated, did not adhere to gelatin-coated plates and were not viable after 3 days in culture (<1:1000). All P2 and P4 adherent cells showed contractility (Supplementary Fig. 5b; Supplementary Movies 3, 4) whereas adult cells adhered to gelatin-coated plates, survived for a few days and expressed troponin T, but failed to contract (Supplementary Fig 5c). The non-organized pattern of troponin T observed on contractile HSA^+^ CMs in culture reflects their immaturity (Fig. 4f) and is a feature also observed during development (Fig. 4e).

**Fig. 5.**
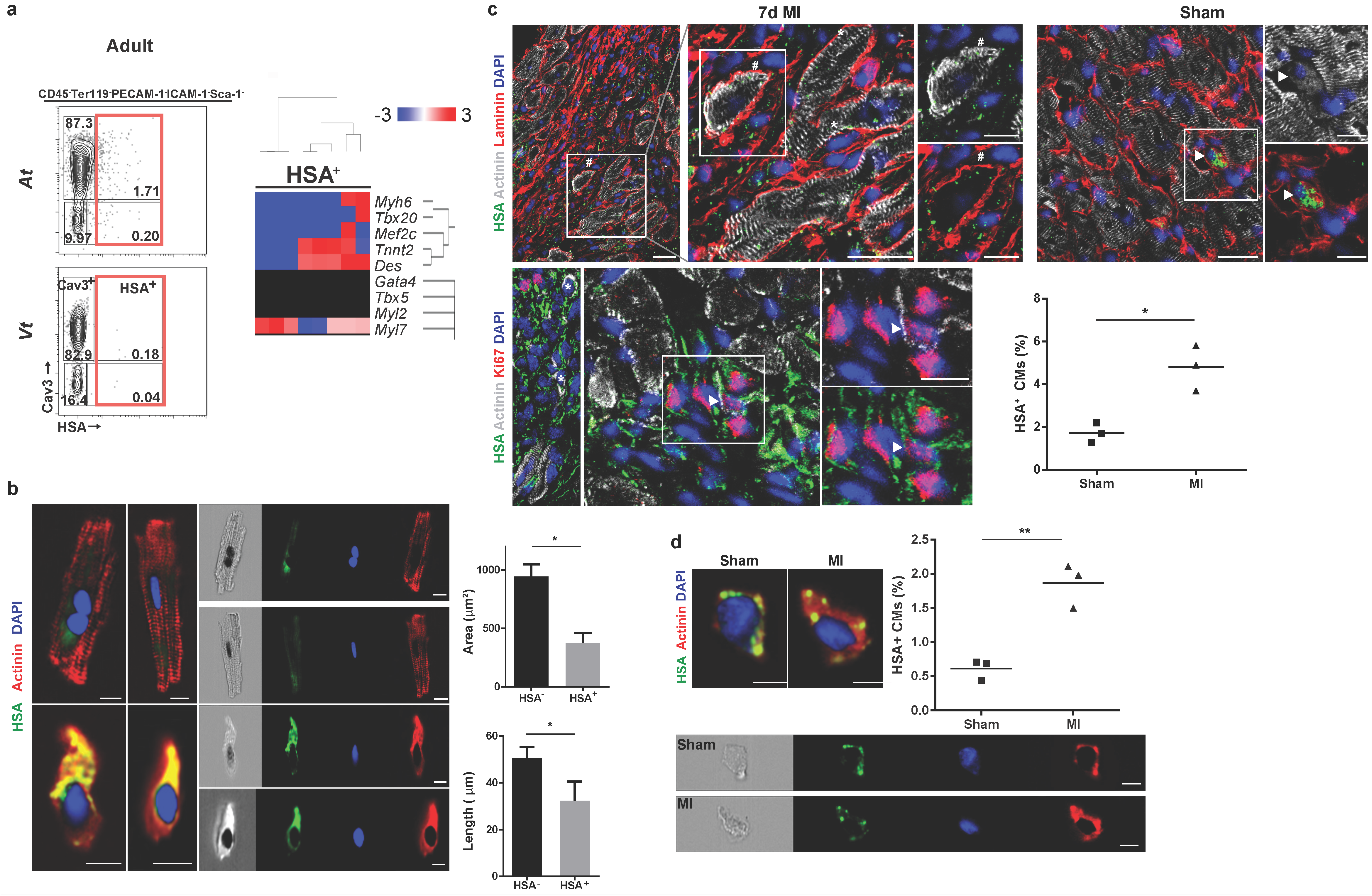
Immature cardiomyocytes in the adult heart respond to injury. **a.** Flow cytometry frequency (top) and gene expression of adult HSA^+^ single cells (bottom) (n=2). **b.** Image flow cytometry of HSA^-^ and HSA^+^ adult CMs. Cells in the top panels show low levels of autofluorescence in the green channel. Histogram bar graphs show length and area of HSA^+^ and HSA^-^CMs (n=3, 9415 cells). **c.** Immunofluorescence of the peri-infarcted myocardium of MI (left panels) or sham-operated animals (right panel). HSA^+^CMs (arrowhead), small round (*) or large striated HSA^+^CMs (#) Ki67^+^HSA^+^CMs (arrowhead). Frequency of HSA^+^CMs (Actinin^+^) per heart after MI (49 sections across 3 hearts) and in sham-operated animals (30 sections across 3 hearts). * p<0.05. Scale bar: 20 µm, insets 10 µm. **d.** Imaging flow cytometry of CMs in MI and sham-operated hearts. Frequency of HSA^+^CMs (Actinin^+^) isolated from sham-operated (n=3, 43461 cells analyzed) and MI hearts (n=3, 35189 cells analyzed). Scale bar: 10 µm. ** p<0.005. CMs | Cardiomyocytes; MI | Myocardial infarction; *At* | Atria; *Vt* | Ventricles.

To show that immature CMs have the capacity to integrate cardiac tissue *in vivo* we used a previously described experimental model^26^, where cardiac cells are transplanted in an ectopic heart tissue previously implanted in the ear-pinna of adult mice (Supplementary Fig. 6a). Viable functional implants were identified by autonomous beating ascertained by visual inspection and by the presence of Ki67^+^ mitotic cells (Supplementary Fig 6b). Seven days later, we transplanted either HSA^+^ CMs or cardiac stromal cells from Ubiquitin-GFP E15.5 embryos, as control (Supplementary Fig. 6a). One week after transplantation, the implants injected with HSA^+^GFP^+^ CMs displayed regions of GFP^+^ cells with the characteristic striated pattern of myocytes, that also co-expressed troponin T (Fig. 4g). Together, these results demonstrated the *in vivo* ability of immature HSA^+^ CMs to engraft cardiac tissue and maintain viability for up to one week.

**Fig. 6.**
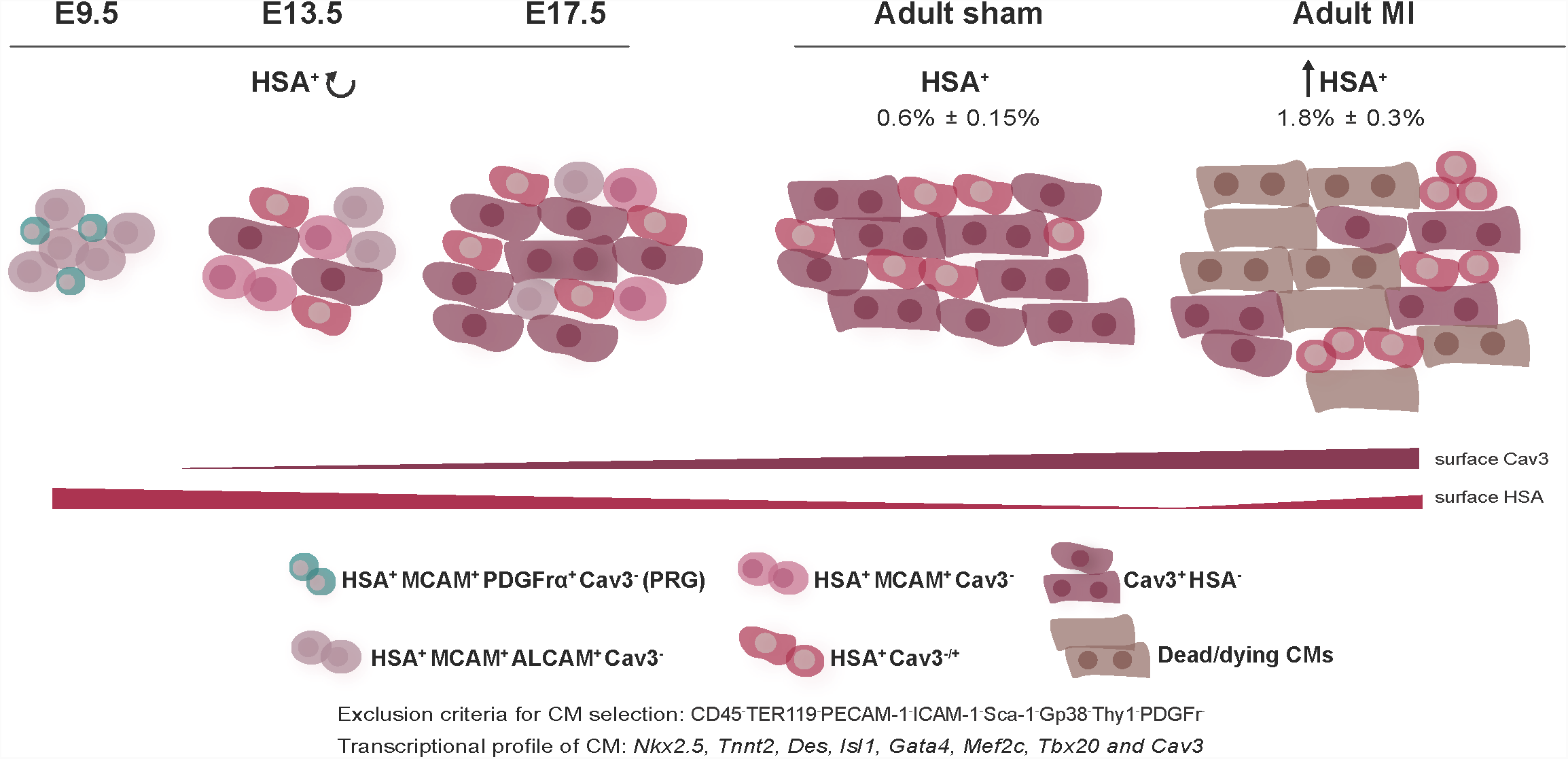
HSA is as a transversal marker of immature CMs during life. The combined use of surface marker analysis with single cell transcriptional profiling allowed identifying different subsets of CMs (expressing *Nkx2.5, TnnT2, Des, Isl1, Gata4, Mef2c* and *Tbx20*) in the murine heart. Less mature CMs (Tnnt^+^Cav3^-^) express HSA, MCAM and ALCAM and progressively loose these markers, maintaining the expression of HSA. Cav3 transcripts are, however, detected in all CMs subsets. A subset of CM PRGs (HSA^+^ALCAM^+^PDGFrα^+^) was found at E9.5 and persisted in the atria of E13.5 and E17.5 hearts (not represented). HSA^+^ CMs dramatically decrease around birth, coinciding with an increase in quiescent and binucleated Cav3^+^ cells, firstly detected at E17.5. The adult myocardium is composed by two main CMs subsets: mononucleated HSA^+^Cav3^+^ and mono- and bi-nucleated Cav3^+^ CMs. Upon MI, HSA^+^ CMs frequency increases and a small percentage of these is proliferating. Throughout cardiac maturation, proliferative CMs (P) were only detected within the HSA^+^ compartment. Detailed information can be found in the text.

Overall, our results identify distinct stages of CM maturation along development based on surface marker expression. We defined two major CMs subsets: an immature subset of mononucleated cells with proliferative capacity (HSA^+^) that can give rise to functional CMs both *in vitro* and *in vivo*; and a mature fraction (Cav3^+^) with increased sarcomere complexity, binucleation and decreased proliferative capacity.

### Immature CMs persist in the adulthood and increase after injury

To determine whether immature HSA^+^ CMs persist in adulthood, we analyzed adult heart cell suspensions with the antibody panel defined above. CM specific transcripts (*Tnnt2, Myl7* and *Myh6*) were present in HSA^+^ cells, more frequent in the *At (At*: 1.7% and *Vt*: 0.18%, Fig. 5a), but also in HSA^-^ cells expressing surface Cav3 (Fig. 5a; Supplementary Fig. 7b). Imaging flow cytometry analysis further confirmed a discrete subset of HSA^+^Actinin^+^CMs in adult hearts (0.6% ± 0.33%, n=3) and showed HSA expression restricted to mononucleated CMs smaller (in area and length) than HSA^-^ CMs (Fig. 5b). The HSA^+^ subset was considered as the most immature adult CMs because they shared cytological and phenotypic properties with embryonic CMs, although the majority also co-expressed the mature marker Cav3. Two stromal populations were also discriminated in the adult: i) a PDGFrα^+^Sca-1^+^Thy1^low^ subset of interstitial FBs compatible with a previously described stromal population^27^ and ii) ICAM-1^+^Gp38^+^Thy1^+^ cells, expressing *Gata4, Tek, Dcn, Twist1* and *Tbx18* and located in the sub-epicardial region (Supplementary Fig. 7b, c). c-kit expression previously associated with CM PRGs was exclusively found in adult PECAM-1^+^ECs and transcripts were also detected in ICAM-1^+^ sub-EpiCs (Supplementary Fig. 7a-b).

The detection of an immature CM subset in the adult prompted us to investigate its frequency in the diseased heart. We found HSA expression largely circumscribed to the non-CM compartment in sham-operated hearts, although rare HSA^+^Actinin^+^CMs were also detected (outlined by laminin expression, Fig. 5c). Seven days after MI, and in spite of HSA expression associated to the upsurge of the hematopoietic and endothelial cells (Supplementary Fig. 7d), we detected a three-fold increase in the frequency of HSA^+^ small round and large striated mononucleated CMs (#) (4.7% ± 3.6%, *) compared to sham-operated (1.7% ±1.1%, Fig. 5c). HSA^+^CMs were found in the peri-infarcted region as shown in the lower magnification image (Fig. 5c). Moreover, a small percentage of HSA^+^CMs (i.e. approximately one per section) were in cycle (Ki67 expression) after MI. A similar increase in HSA^+^CMs was evidenced after MI by imaging flow cytometry analysis of pre-fixed cells (1.8% ± 0.3%), when compared to sham-operated hearts (0.6% ± 0.15%, Fig. 5d).

Our results show that the cell surface signatures defined for the embryonic heart are suitable to isolate an affiliated adult population. Importantly, we identified a subset of HSA-expressing CMs, with less differentiated features than the majority of adult CMs, that cycles and increases in frequency after MI.

## Discussion

The mouse heart is able to regenerate during the first days of life by proliferation of pre-existing CMs. This capacity is lost after one week of postnatal life, time from which wounded tissue is replaced by a non-functional scar^28^. The loss of CM mitotic activity has been attributed to the binucleation process and the complete maturation of CMs that occur after birth^10,29^. However, recent reports showed that CM replacement and cell division could occur, although at low rate^21,22^, raising the possibility that a subset of CMs in the adult might undergo mitosis. The low frequency of dividing CMs requires the identification of specific markers to isolate and characterize these rare cells. We show here that HSA marks the proliferative neonatal CM compartment that persists in the adult, retains some of their embryonic features and expands after MI.

Expression of *Isl1, Gata4, Mef2c* and *Tbx20* is important for CM commitment and initial stages of differentiation^30^-^32^ and accordingly were used as indicative of the CM lineage. Consistent with previous reports^27^, we found some of these transcripts also expressed during the development of stromal cells (SMCs and FBs) indicating that, in the heart, they participate in the development of different lineages. These findings demonstrate that CMs can only be unambiguously identified by the combined expression of transcription factors, transcripts codifying structural and contractile proteins and by the absence of stromal or ECs-associated markers.

Several studies identified CM PRGs based on the expression of Sca-1 and c-kit^14^-^16^ cell-surface proteins, which in our work were not found within the CM compartment. c-kit was only expressed in ECs (PECAM-1^+^), as recently shown also by others^34^, and in Thy1^+^PDGFrα^+^ FBs. Sca-1 expression was detected in a fraction of ECs and in a population of cells in the AV canal in the fetal heart and in the interstitium of adult myocardium^15^. Their spatial pattern and transcriptional profile are compatible with a FB lineage affiliation, supported by the description of Sca-1-expressing cells exhibiting a paracrine role in angiogenic stimulation, after MI^15,35^. At E9.5 we found a HSA, ALCAM and PDGFrα expressing population that was highly proliferative and expressed *Nkx2-5* and *Tnnt2* together with *Isl1* and *Tbx5* suggesting they represent CM PRGs^36,37^. These cells sharply decrease in frequency, they are only detected in the *At* after E13.5 of development and become undetectable after birth, a finding that is not compatible with the persistence of CM PRGs in the postnatal heart.

We identified HSA, so far never associated to heart development or maturation, as a transversal marker of immature CMs throughout life. Our analysis showed a continuum in CM maturation, which is an asynchronous process that starts during development and can be prospectively identified by the expression of distinct surface markers (Fig. 6). Immature CMs can be identified by a unique phenotype, i.e. HSA^+^MCAM^+^ALCAM^+^Cav3^-^, they progressively lose ALCAM and MCAM expression to become HSA^+^ only. HSA^+^ CM decrease in frequency between E17.5 and P7, but are the only CM subset that actively proliferates up to P7, they have spontaneously contractile properties in culture and are consistently mononucleated. Isolated from E15 hearts HSA^+^ CM engrafted cardiac tissue transplanted in the ear pinna of adult mice.

The first signs of CM binucleation are observed around E17.5 of development in Cav3^+^ CMs, but never in immature CMs (Tnnt^+^Cav3^-^) that decrease in frequency, coinciding with an increase in non-cycling and binucleated Cav3^+^ CMs that compose the myocardium in adulthood. These results are in agreement with alterations endured by CMs during the first week of life, which encompass a transition from hyperplasia to hypertrophy and terminal differentiation^9,10^.

A subset (<1%) of adult CMs displayed HSA and remained mononucleated, thus resembling embryonic CM, despite expressing Cav3 at the cell surface. Our findings are in line with previous reports showing low rate of cell division (0.76%^21^ and 0.3%-1%^33^ per year) in small, mononucleated and diploid CMs, in the adult heart.

Foreseeing its therapeutic relevance, we tested whether HSA^+^ immature CMs could respond to a pathological challenge. Interestingly, we observed an upsurge in the frequency of adult mononucleated HSA^+^ CMs following 7 days of MI (from 0.6% to 1.8%). This relative increase on HSA^+^ CMs can be explained by proliferation, detected at low frequency in our analysis, by increased resistance of HSA^+^CMs to hypoxia, by re-expression of HSA or by any combination of the above. In the developing heart, low oxygen tension is found in the compact myocardial layer^38^ precisely where immature HSA^+^CMs were found in this study. Likewise, an adult CM subset protected from oxidative stress in hypoxic niches and exhibiting low proliferative activity upon injury has been recently identified. Similar to HSA^+^CMs, these cells were mononucleated, small sized and represented around 1% of the adult myocardium^33^. It is thus tempting to hypothesize that the cells described in both studies correspond to the same population.

Although HSA^+^ immature CMs do not proliferate sufficiently to regenerate the myocardium, they might account for the low CM turnover rate previously described in the adult^21,22,39^ and be more amenable than binucleated CMs to respond to mitotic stimuli. Importantly, using the strategy herein described, CMs at different maturation stages can now be prospectively isolated as viable cells from the adult heart, enabling further mechanistic studies.

## METHODS

### Mice

C57BL/6 mice (Charles River) 6 - 8 week-old or timed pregnant females were used. Timed-pregnancies were generated after overnight mating. The following morning, females with vaginal plug were considered to be at E0.5.

All animal manipulations at i3S were approved by the Animal Ethics Committee and Direcção-Geral de Veterinária – DGAV; at Pasteur Institute according to the ethic charter approved by French Agriculture ministry and to the European Parliament Directive 2010/63/EU at both institutes.

### Mouse model of Myocardial Infarction

MI was experimentally induced by permanent ligation of the left coronary artery as previously described^40^, and samples analysed 7 days after injury.

### Transplantation of Ubiquitin–GFP cells in embryonic cardiac implants (ear-pinna model)

As shown in Supplementary Fig. 6a, E15.5 cardiac *Vt* from wild-type (WT) embryos were dissected and grafted in the ear-pinna of a recipient adult WT mice, under anesthesia, as previously described^26^. Seven days later, E15.5 hearts from Ubiquitin–GFP mice were dissociated and HSA+ immature CMs or PDGFrα+ fibroblasts (stromal cells) were sorted and directly injected into visible beating implants (10 000 cells per implant). One week later, the implants were collected and tissue was processed for immunofluorescence as described below.

### Isolation of Live Cardiac Cells

Embryonic hearts were collected under a stereomicroscope and the three anatomic heart structures (*At, GV-AVJ* and *Vt*) were micro-dissected. Heart tissue was minced into 1 mm^3^ fragments and incubated during 15 minutes at 37°C in the enzymatic solution: for E13.5 and E17.5 hearts, 0.2 mg/mL collagenase (Sigma) in Hank’s Balanced Salt Solution with calcium and magnesium (HBSS+/+, Invitrogen); for E9.5 hearts, 0.1 mg/mL collagenase in HBSS+/+; and adult heart, 0.2 mg/mL collagenase with 60 U/mL DNase I (Roche). At the end of each round of digestion, tissue fragments were re-suspended using a P1000 pipette (approximately 20 times). The remaining tissue was let to sediment and the supernatant was collected in a tube containing the same volume of 10% FCS (Gibco)-HBSS+/+ and kept on ice, while the digestion protocol continued. Digestion was repeated until no macroscopic tissue was detected. After digestion, cell suspensions were centrifuged 10 minutes, 290 g at 4°C, re-suspended in 1% FCS-HBSS+/+ and filtered with a 70 µm mesh strainer (Fisher).

### Isolation of Fixed Cardiomyocytes

Fixed cardiomyocytes were isolated as described by Mollova and colleagues^41^ with some alterations. E13.5, E17.5, P7, Adult and injured (MI or Sham-operated) hearts were collected, washed in PBS (to remove blood), minced into 2 mm^3^ pieces, flash frozen in liquid nitrogen and stored at -80°C. For cell isolation, tissue pieces were fixed in 4% paraformaldehyde at room temperature for 2 hours, washed in PBS for 5 minutes and digested with 3 mg/ml collagenase type II (Worthington) in HBSS on a rotator at 37°C until no macroscopic tissue was detected. Enzyme activity was blocked with 10% FBS-HBSS (Gibco). Cell suspensions were filtered through a 100 µm cell-strainer.

### Flow Cytometry, Cell Sorting and Imaging Flow Cytometry

Heart cell suspensions were stained with (conjugated or non-conjugated) antibodies (20 minutes, 4°C in the dark) followed by incubation with conjugated streptavidin (10 minutes, 4°C in the dark). Whenever using a non-conjugated antibody, a sequential incubation with a secondary antibody was performed 15 minutes 4°C in the dark (see Supplementary Table 2 for the antibodies list). Propidium iodide (PI, 1µg/ml) was used to exclude dead cells. Intracellular proteins detection was performed after surface staining, fixation and permeabilization with the Foxp3/Transcription Factor Staining Buffer Set (eBioscience). DAPI was used to stain DNA in fixed cells (5 minutes at 4°C). Flow cytometry data was acquired in a BD FACSCanto™ II and in a Sony SP6800 analyser (Sony) and analysed with the FlowJo v10.0.8 (Treestar), Kaluza 1.5 (Beckman Coulter) or R v3.2.4 software. Cells were sorted in a BD FACSAria™ III directly into 96-well plates loaded with RT-STA Reaction mix (CellsDirect™ One-Step qRTPCR Kit, Invitrogen, according to the manufacturer’s procedures) and 0.2x specific TaqMan^®^ Assay mix (see Supplementary Table 4 for the TaqMan^**®**^ assays list). For single cell sorting a control-well with 20 cells was always used. Index sorting tool on BD FACSDiva™ v8.0.1 software (BD Bioscience) was activated to track and record the fluorescence data for each parameter of each individual cell collected in a precise position of the 96-well plate. This tool allowed the post-sorting correlation of the levels of surface protein expression and the transcriptional profile (Supplementary Fig. 2c). Data were analysed with the FlowJo v10.0.8 software.

Fixed cardiomyocytes were re-suspended in BD Cytofix/Cytoperm Fixation/Permeabilization Kit (BD Biosciences), permeabilized in BD Perm/Wash™ buffer for 15 minutes, incubated with primary antibodies for 2 hours at 4°C and with Alexa Fluor-conjugated secondary antibodies for 30 minutes at 4°C. Prior to acquisition on ImageStream, nuclei were stained with 20 µM DRAQ5 (Biostatus) and filtered with 100 µm mesh. Data acquisition was performed using an ImageStreamX cytometer (Amnis). Files were collected with a cell classifier applied to the bright-field (BF) channel to capture events larger than 20µm and included BF, FITC, PECy7 and DRAQ5 images. At least 30000 cells were analyzed for each sample and all images were captured with the 40x objective. Data analysis was performed with IDEAS software (v6.0, Amnis). For each sample only intact cardiomyocytes, selected based on Actinin and DRAQ5 signal intensity, were considered for subsequent analysis. For the morphometric analysis we applied a morphology mask to the BF channel and for assessment of the number of nuclei/cell we used the DRAQ5 images.

### Culture of HSA+ cells and Live-cell imaging

E15.5, P2, P4 and adult cardiac cells were isolated as above and HSA^+^ cells were sorted following the gating strategy in Supplementary Fig. 5a-b. For the neonatal and adult cells, 20 mM 2,3-Butanedione monoxime (BDM, Sigma) was added throughout the isolation procedure^42^. HSA^+^ cells were plated in 0.1% collagen (Life Technologies) for E15.5 or fibronectin/gelatin for postnatal cells, coated ibidi plates and cultured for one week in high glucose Iscove’s Modified Dulbecco’s media (Gibco, Life Technologily) supplemented with 20%FBS, 1X penicillin/streptomycin (Gibco, Life Technologies), 1X L-glutamine (Gibco, Life Technologies), 50 µg/mL µg/ml ascorbic acid, and 1.5×10-4M 1-thioglycerol (Sigma-Aldrich), as previously described^4^. Adult cardiac cells were incubated at 37°C in 3%O2. Live-cell imaging was performed on a temperature-controlled Zeiss Axiovert 200M microscope equipped with a CoolSnap HQ (Roper) camera. Sample position was controlled by a X-Y motorized stage and images were acquired every 15 min using an A-Plan 20x/0.30 objective for 48 hours.

### Histological Processing and Immunofluorescence Staining

Embryonic and adult (MI and sham-operated) hearts were fixed in 0.2% paraformaldehyde (Merk) overnight at 4°C, dehydrated in a sucrose gradient (4% followed of 15%), embedded in gelatine and frozen. Tissue cryo-sections (4 µm thick) were blocked with either 4% FBS-1% BSA blocking solution or Vector M.O.M. ™ basic kit (Vector Laboratories), depending on the specific conditions detailed on Supplementary Table 3. Tissue sections were incubated with primary antibodies overnight at 4°C, followed by 1-hour incubation with Alexa Fluor-conjugated secondary antibodies (Invitrogen, see Supplementary Table 3 for the antibodies list). Slides were mounted and nuclei counterstained with aqueous mounting medium with DAPI (Vector Labs). Representative high-resolution images were acquired for each heart structure (*At, GV-AVJ* and *Vt*) at 40x magnification in a confocal microscope (Leica SP5II, Leica). Whole heart acquisitions were obtained using the high-content imaging system (IN Cell Analyzer 2000, GE Healthcare).

Isolated fixed cardiomyocytes were re-suspended in 10% FBS-PBS and spun onto superfrost slides in a cytocentrifuge (Shandon). Cytospins were incubated with primary antibodies overnight at 4°C, followed by 1-hour incubation with Alexa Fluor-conjugated secondary antibodies. Acquired images were edited and quantified using the Image J software v1.51d software.

### Gene Expression Analysis

Sorted cells in RT-STA Reaction mix from the CellsDirectTM One-Step qRT-PCR Kit (Life Technologies) were kept at -80°C at least overnight before reverse transcription and specific target pre-amplification (20 cycles for single cells and 18 cycles for 20 cells). Pre-amplified samples were subjected to qPCR (see Supplementary Table 4 for Taqman® assays list) as previously described^43^.

### Bioinformatic Analysis

Flow cytometry data analysis was performed in FCS files of live CD45^-^Ter119^-^CD31^-^ cell fraction using R package flowCore from R v3.2.4 revised (2016-03-16 r70336) and the interface R Studio v0.99.467^44^. Subsequently gating, as described in Fig. 1 and in Supplementary Fig. 1, was used to define each population. Map clustering of the flow cytometry data was performed using custom R scripts from R package t-SNE to dimensionality reduction – t-SNE^45^ and Bioconductor.org package flowSOM to visualize Self-Organizing and Minimal Spanning Trees (Spanning Trees)^46,47^.

Gene expression raw data (BioMark^™^, Fluidigm) of sorted cells at the population level was normalized with HPRT, and data is presented in 2^(-ΔCt). Single cell gene expression analysis was performed in cells that expressed at least one of three housekeeping genes (*Hprt, Gapdh*, or *Actb*) and Ct values were used to the following analysis. A Ct value of 21 was the maximum value considered as expressed gene and the background (i.e. non-detected) Ct value was 38. qPCR data was processed with the ^©^QLUCORE (Qlucore AB 2008 – 2015, Lund, Sweden) software and was displayed in uncentered Pearson’s correlation unsupervised hierarchical clustering and Principal component analysis (PCA) either for surface phenotype or transcripts.

### Statistical Analysis

All results are shown as mean ± standard deviation (SD). Statistical significance was determined using the t-Student test, except when comparing the frequency of HSA^+^ CMs form E13.5 to P7 (one-way ANOVA followed by Tukey Test). The statistical analysis of the data was performed using SigmaPlot software (*p*<0.05 was considered statistically significant) or ^©^QLUCORE (Qlucore AB 2008 – 2015, Lund, Sweden) software for the multidimensional analysis of multiplex qPCR (two-way ANOVA, *p*=0.007, *q*=0.01).

## Acknowledgments

We thank the members of P.P.O. and A.C. laboratories for fruitful discussions. We are grateful to the i3S Animal Facility, Advanced Flow Cytometry, Unit b.IMAGE, Advanced Light Microscopy Unit and to the Pasteur Institute Flow Cytometry Core facility, including Sony SP6800 implementation (K. Futamura and C. Ait-Mansour), D. Montarra’s and S. Meilhac’s lab, P. Vieira, A. Bandeira, H. Maiato and specially to P. Pereira for critical reading of the manuscript; M. Rujano, Z. Garcia, D. Sassoon, S. Tajbakhsh; P. Bousso for the ubiquitin-GFP reporter mice; S. Chea and V. Rouilly for help in the analysis of single cell transcriptional data.

This work was financed by European Structural and Investment Funds (ESIF), under Lisbon Portugal Regional Operational Program and National Funds through FCT-Foundation for Science and Technology under project POCI-01-0145-FEDER-016385 to PPO; by Pasteur Institute, INSERM, ANR (grant Twothyme), REVIVE Future Investment Program and Pasteur-Weizmann Foundation through grants to AC. MV (SFRH/BD/74218/2010) and TPR (SFRH/BPD/80588/2011) were supported by FCT, and PPO was recipient of an invited scientist grant by Institut Pasteur, Paris, France

## Auth or contributions

M.V., T.P.R., D.S.N., A.C. and P.P.O. designed the project; M.V., T.P.R., D.S.N., O.B.-D. and A.C. performed experiments and analyzed data; B.D. contributed new analytic tools; and M.V., T.P.R., D.S.N., A.C. and P.P.O. wrote the paper.

## Additional information

### Supplementary Information

Supplementary Information accompanies this paper at

### Reprints and permission

Information is available online at Correspondence and requests for materials should be addressed to A.C. and to P.P.O.

