## Supplemental file for "HSA^+^ immature cardiomyocytes persist in the adult heart and expand after ischemic injury"

#### **Supplementary Information Inventory:**

##### **SUPPLEMENTARY TABLES:**

**Supplementary Table 1. Summary of the characteristics of the newly defined E17.5 cardiac lineages**

**Supplementary Table 2. List of antibodies for flow cytometry**

**Supplementary Table 3. List of antibodies for immunofluorescence**

**Supplementary Table 4. List of TaqMan® Assays**

##### **SUPPLEMENTARY FIGURES:**

**Supplementary Fig. 1. Gating strategy used to define the cardiac cell populations**

**Supplementary Fig. 2. Single cell transcriptional profiles of cardiac cell types at E17.5**

**Supplementary Fig. 3. Surface phenotype progression of the HSA+ CMs during heart morphogenesis and their cell cycle status**

**Supplementary Fig. 4. Detailed analysis of the two subsets of cardiomyocytes for doublet discrimination and Tnnt expression**

**Supplementary Fig. 5. Purity analysis and quantification of the HSA+ CM cultures**

**Supplementary Fig. 6. Ear-pinna transplantation experiments**

**Supplementary Fig. 7. Adult cardiac stromal populations**

##### **SUPPLEMENTARY MOVIES:**

**Supplementary Movie 1. E15.5 HSA+ cardiomyocyte dividing**

**Supplementary Movie 2. E15.5 HSA+ cardiomyocyte beating**

**Supplementary Movie 3. P2 HSA+ cardiomyocyte beating**

**Supplementary Movie 4. P4 HSA+ cardiomyocyte beating**

### SUPPLEMENTARY TABLES

Supplementary Table 1. Summary of the properties of the newly defined E17.5 cardiac lineages

| E17.5 |  |  |  |  |  |
| --- | --- | --- | --- | --- | --- |
| Cluster | Genes | Cell Type | Location |  | Cell Surface Phenotype |
|  |  |  | Anatomical | Histological |  |
| Ia | *Nkx2-5*<sup>***</sup>,<br>*Tnnt2*<sup>***</sup>, *Des*<sup>***</sup>,<br>*Myl7*, *Myh6* | At CMs | *At* | Widespread in myocardium | **HSA<sup>+</sup>MCAM<sup>+</sup>**<br>**ALCAM<sup>+</sup>** |
| Ib | *Nkx2-5*<sup>***</sup>,<br>*Tnnt2*<sup>***</sup>, *Des*<sup>***</sup>,<br>*Myl2*, *Myh7* | Vt CMs | *Vt* | Small *foci* in myocardium | **HSA<sup>+</sup>MCAM<sup>+</sup>** |
| II | *Acta2*<sup>***</sup>,<br>*Myh11*<sup>***</sup> | SMCs | GV-AVJ | Great vessels wall | **ALCAM<sup>+</sup>** |
| III | *Kdr*<sup>***</sup>, *Flt1*<sup>***</sup>,<br>*Tek*<sup>***</sup> | ECs | *At*, GV-AVJ,<br>*Vt* | Endothelium and endocardium | **PECAM-1<sup>+</sup>** |
| IV | *Tbx18*<sup>***</sup>, *Wt1*<sup>***</sup>,<br>*S100a4*<sup>***</sup>,<br>*Gja1*<sup>***</sup> | EPDCs | *At*, GV-AVJ,<br>*Vt* | Sub-EpiC zone | **ICAM-1<sup>+</sup>** |
| V | *Twist1*<sup>***</sup>,<br>*Tbx3*<sup>***</sup>, *Vcan*<sup>***</sup>,<br>*Fap*<sup>***</sup>, *Vim*<sup>***</sup>,<br>*Isl1*<sup>***</sup> | GV-AVJ and EndoC cushions FBs | GV-AVJ | EndoC cushions | **HSA<sup>+</sup>** |
|  |  |  |  | Great vessels insertion | **Sca-1<sup>+</sup>** |
| VI | *Tbx20*, *Snai2*,<br>*Vim*, *Fn*, *Fap* | SMCs | GV-AVJ, *Vt* | - | **MCAM<sup>+</sup>** |
| VII | *Ddr2*, *Col3a1*,<br>*Dcn*, *Postn*, *Fn1*,<br>*Vim*, *Snai2*, *Tbx20* | EpiCs | *At*, GV-AVJ and *Vt* | Epicardium | **Gp38<sup>+</sup>** |
|  |  | FBs | GV-AVJ, *Vt* | - | **Thy1<sup>+</sup>** |
|  |  |  | *At*, GV-AVJ, *Vt* | Interstitium | **PDGFrα<sup>+</sup>** |

One way-ANOVA \*\*\* $p = 0.000435$; $q = 0.0022$

**Supplementary Table 2. List of antibodies for flow cytometry**

| Protein |  | Clone | Fluorochrome | Source | Reference |
| --- | --- | --- | --- | --- | --- |
| Caveolin 3 | Cav3 | - | - | BD Biosciences | 610420 |
| CD146 | MCAM | ME-9F1 | FITC | Miltenyi biotec | 130-102-230 |
| CD166 | ALCAM | eBioALC48 | APC | eBioscience | 17-1661-82 |
| CD24 | HSA | M1/69 | PECy7 | BD Bioscience | 560536 |
| CD31 | PECAM-1 | MEC13.3 | Alexa Fluor® 647 | BioLegend | 102516 |
| CD31 | PECAM-1 | MEC13.3 | PE | BD Bioscience | 553373 |
| CD31 | PECAM-1 | MEC13.3 | BV421 | BD Bioscience | 562939 |
| CD45 | - | 30-F11 | PE | BioLegend | 103106 |
| CD45 | - | 30-F11 | B | BioLegend | 103104 |
| CD54 | ICAM-1 | YN1/1.7.4 | PB | BioLegend | 116116 |
| CD54 | ICAM-1 | 3E2 | B | BD Bioscience | 553251 |
| CD90.2 | Thy1 | 30-H12 | FITC | BD Bioscience | 553013 |
| CD90.2 | Thy1 | 30-H12 | PE | BD Bioscience | 553014 |
| CD90.2 | Thy1 | 53-2.1 | BV605 | BD Bioscience | 563008 |
| c-Kit | - | 2B8 | APCCy7 | BioLegend | 105825 |
| Gp38 | - | eBio8.1.1 (8.1.1) | eFluor® 660 | eBioscience | 50-5381-82 |
| Ki67 | - | MOPC-21 | FITC | BD Bioscience | 556026 |
| PDGFra | - | APA5 | PE | BioLegend | 135905 |
| Sca-1 | - | D7 | PECy5 | BioLegend | 108110 |
| Ter119 | - | TER-119 | PE | BioLegend | 116208 |
| Ter119 | - | TER-119 | B | BD Bioscience | 553672 |
| TroponinT | Tnnt | 13-11 | - | Thermo Scientific | MS-295-P0 |
| Conjugated-secondary antibodies and – streptavidin |  |  |  |  |  |
| Donkey Anti-Mouse IgG |  | - | Cy3 | Jackson ImmunoResearch Laboratories | 715-167-003 |
| SAV |  | - | APCCy7 | BioLegend | 405208 |
| SAV |  | - | BV421 | BioLegend | 405225 |

**Supplementary Table 3. List of antibodies for immunofluorescence**

| Primary antibodies |  |  |  |  |  |
| --- | --- | --- | --- | --- | --- |
| Protein |  | Isotype | Conditions | Source | Reference |
| Actinin | - | Mouse IgG | 1:300 | Sigma | A7811 |
| α-Smooth Muscle Actin | SMA | Mouse IgG | 1:300 | Sigma | A5228 |
| Caveolin 3 |  | Mouse IgG | 1:150 | BD Bioscience | 610420 |
| CD166 | ALCAM | Rat IgG | 1:50 | eBioscience | 17-1661-82 |
| CD24 | HSA | Rat IgG | 1:150 | eBioscience | 14-0242-81 |
| CD31 | PECAM-1 | Goat IgG | 1:250 | SCBT | sc-1506 |
| CD54 | ICAM-1 | Rat IgG | 1:300 | eBioscience | 14-0541 |
| Gp38 | - | Hm IgG | 1:300 | Novus | NB600-1015SS |
| Ki67 | - | Rabbit IgG |  | Abcam | ab15580 |
| Laminin | - | Rabbit IgG | 1:500 | Sigma | L9393 |
| PDGFRα | - | Goat IgG | 1:500 | R&D Systems | AF1062 |
| Sca-1 | - | Rat IgG |  | BD Pharmingen | 553333 |
| Secondary antibodies |  |  |  |  |  |
| Antibody |  | Fluorochrome |  | Source | Reference |
| Donkey Anti-Mouse IgG |  | Cy3 |  | Jackson ImmunoResearch Laboratories | 715-167-003 |
| Chick Anti-Mouse IgG |  | 647 |  | Invitrogen | A-21463 |
| Donkey Anti-Rat IgG |  | 488 |  | Invitrogen | A-21208 |
| Donkey Anti-Goat IgG |  | 594 |  | Invitrogen | A-11058 |
| Donkey Anti-Rabbit IgG |  | 568 |  | Invitrogen | A10042 |
| SAV |  | PECy7 |  | BioLegend | 405206 |

**Supplementary Table 4. List of Taqman® Assays**

| **Gene** | **Gene Symbol** | **Gene ID** | **Taqman® Assay ID** |
| --- | --- | --- | --- |
| *Actin beta* | *Actb* | 11461 | Mm00607939_s1 |
| *aMHC* | *Myh6* | 17888 | Mm00440359_m1 |
| *bMHC* | *Myh7* | 140781 | Mm00600544_m1 |
| *Caveolin 3* | *Cav3* | 12391 | Mm01182632_m1 |
| *c-Kit* | *Kit* | 16590 | Mm00445212_m1 |
| *Coll I* | *Col1a1* | 12842 | Mm00801666_g1 |
| *Coll III* | *Col3a1* | 12825 | Mm01254476_m1 |
| *Connexin 43* | *Gja1* | 14609 | Mm00439105_m1 |
| *Connexin 40* | *Gja5* | 14613 | Mm01265686_m1 |
| *Connexin 45* | *Gjc1* | 14615 | Mm01253027_m1 |
| *cTnt* | *Tnnt2* | 21956 | Mm01290256_m1 |
| *DDR2* | *Ddr2* | 18214 | Mm00445615_m1 |
| *Decorin* | *Dcn* | 13179 | Mm00514535_m1 |
| *Desmin* | *Des* | 13346 | Mm00802455_m1 |
| *Fap* | *Fap* | 14089 | Mm01329177_m1 |
| *Fibronectin* | <i fn1<="" i=""> | 14268 | Mm01256744_m1 |
| *Flk1* | *Kdr* | 16542 | Mm01222421_m1 |
| *Flt1* | *Flt1* | 14254 | Mm01210866_m1 |
| *Fsp1* | *S100a4* | 20198 | Mm00803371_m1 |
| *Gapdh* | *Gapdh* | 14433 | Mm99999915_g1 |
| *Gata4* | *Gata4* | 14463 | Mm00484689_m1 |
| *HCN4* | *Hcn4* | 330953 | Mm01176086_m1 |
| *HPRT* | *Hprt* | 15452 | Mm01545399_m1 |
| *Islet1* | *Isl1* | 16392 | Mm00517585_m1 |
| *Mef2c* | *Mef2c* | 17260 | Mm01340842_m1 |
| *Mlc2v* | *Myl2* | 17906 | Mm00440384_m1 |
| *Mlc2a* | *Myl7* | 17898 | Mm01183005_g1 |
| *Nfatc1* | *Nfatc1* | 18018 | Mm00479445_m1 |
| *NG2* | *Vcan* | 13003 | Mm01283063_m1 |
| *Nkx2.5* | *Nkx2-5* | 18091 | Mm01309813_s1 |
| *Scleraxis* | *Scx* | 20289 | Mm01205675_m1 |
| *Slug* | *Snai2* | 20583 | Mm00441531_m1 |
| *SMA* | *Acta2* | 11475 | Mm01546133_m1 |
| *sm-MHC* | *Myh11* | 17880 | Mm00443013_m1 |
| *Pax3* | *Pax3* | 18505 | Mm00435491_m1 |
| *Periostin* | *Postn* | 50706 | Mm00450111_m1 |
| *Tbx1* | *Tbx1* | 21380 | Mm00448949_m1 |
| *Tbx2* | *Tbx2* | 21385 | Mm00436915_m1 |
| *Tbx3* | *Tbx3* | 21386 | Mm01195726_m1 |
| *Tbx5* | *Tbx5* | 21388 | Mm00803518_m1 |
| *Tbx18* | *Tbx18* | 76365 | Mm00470177_m1 |
| *Tbx20* | *Tbx20* | 57246 | Mm00451515_m1 |
| *Tcf21* | *Tcf21* | 21412 | Mm00448961_m1 |
| *Tenascin*<br>*C* | *Tnc* | 21923 | Mm00495662_m1 |
| *Tie2* | *Tek* | 21687 | Mm00443243_m1 |
| *Twist1* | *Twist1* | 22160 | Mm04208233_g1 |
| *Vimentin* | *Vim* | 22352 | Mm01333430_m1 |
| *Wnt1* | *Wnt1* | 22408 | Mm01300555_g1 |
| *Wt-1* | *Wt1* | 22431 | Mm01337048_m1 |

### Supplementary Fig. 1

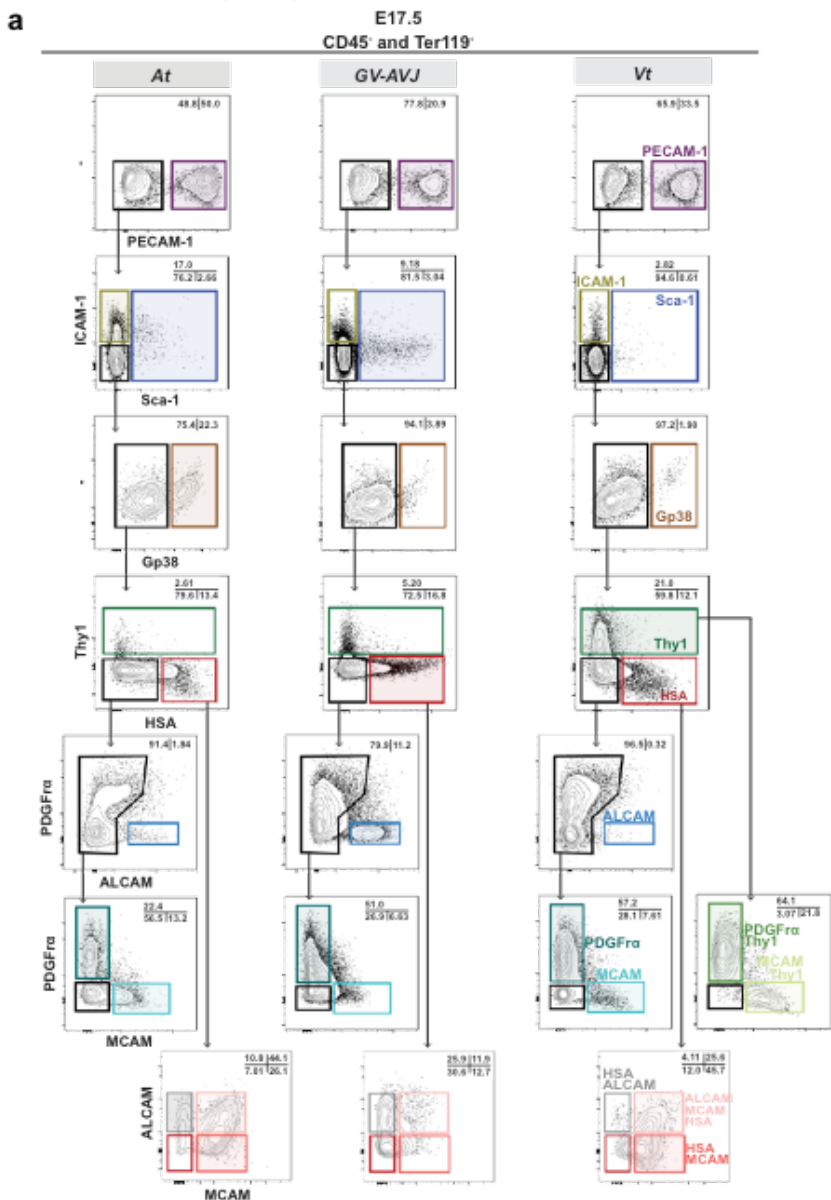

**Supplementary Fig. 1. a.** Gating strategy used to define the cardiac cell populations. A. Single cell suspension from the three heart regions (At | Atria; GV-AVJ | Great Vessels-Atrioventricular Junction; Vt | Ventricles) were analyzed for CD45, Ter119, PECAM-1, ICAM-1, Sca-1, Thy1, HSA, PDGFRα, ALCAM and MCAM markers in a SP6800 Spectral analyzer. Representative contour plots of the indicated surface proteins in the CD45<sup>-</sup>Ter119<sup>-</sup> fractions (the upper plots) and in the subsequent gates indicated by the black arrows are shown and define the gating strategy. Numbers indicate frequencies within the gates. **b.** Listing of the complete surface signature for each cardiac population together with their acronyms and the corresponding color-code.

#### Supplementary Fig. 2

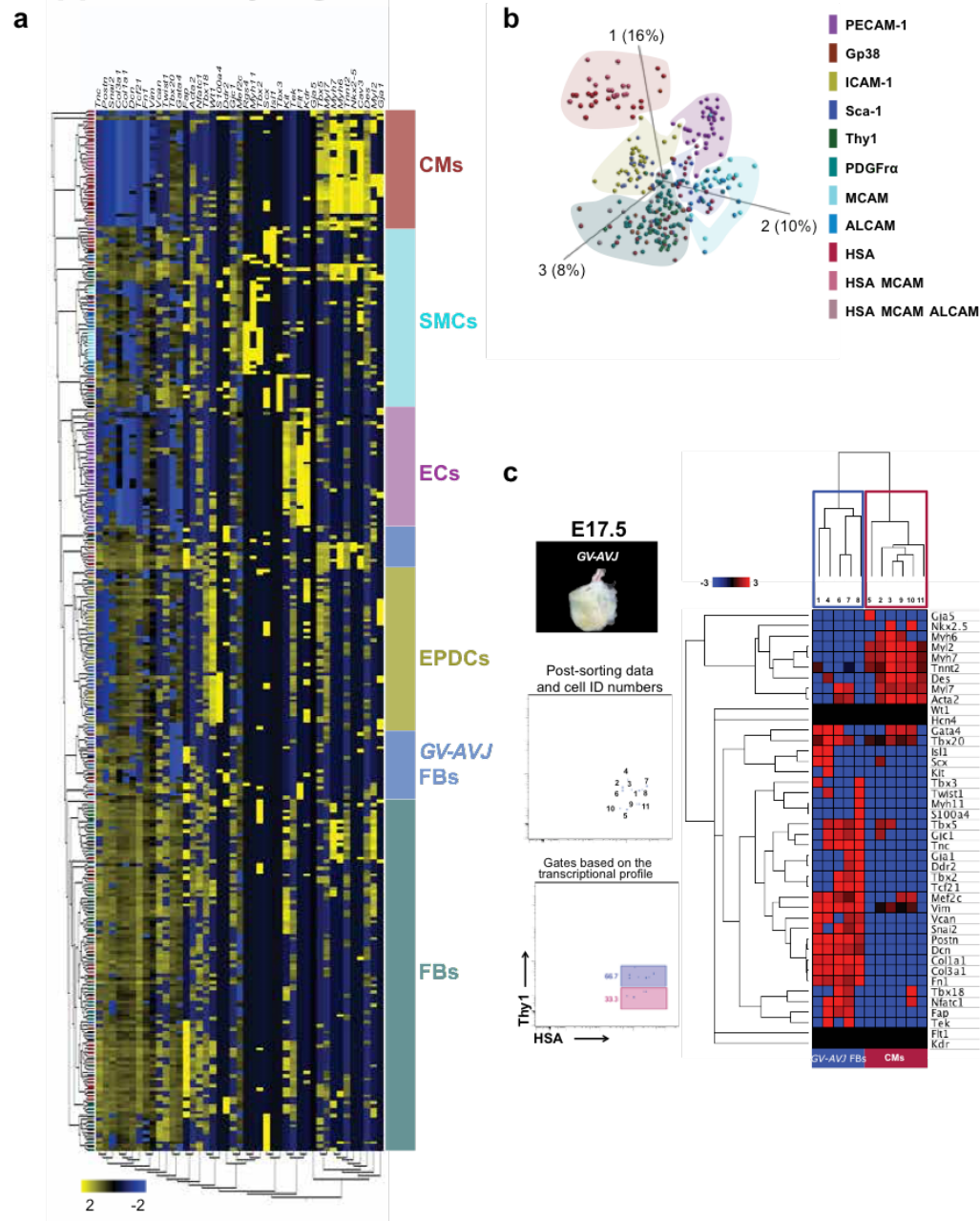

**Supplementary Fig. 2.** Single cell transcriptional profiles of cardiac cell types at E17.5. **a.** Heat Map displays the unsupervised hierarchical clustering analysis of the multiplex single-cell qRT-PCR data of individual cardiac cells (311 analyzed single cells) as in Fig. 2. **b.** PCA graph corresponding to the heat map analysis shown in **a.** **c.** Index-sorting analysis correlates the phenotype of each sorted cell with its transcriptional profile. Macroscopic view of the E17.5 GV-AVJ dissected region showing the recurrent contamination with Vt tissue. Thy1 versus HSA dot plots showing the levels of Thy1 and HSA expression of each sorted cell, to which a number was ascribed. Heat map of the unsupervised hierarchical clustering for the multiplex single cell qRT-PCR performed on the individually sorted cells. Using the index sorting tool, we distinguished by the levels Thy1 expression Vt-derived CM (low) from GV-AVJ HSA<sup>+</sup> fibroblasts (high). GV-AVJ | Great Vessels-Atrioventricular Junction; Vt | Ventricles; CMs | Cardiomyocytes; GV-AVJ FBs | Great Vessels-Atrioventricular Fibroblasts.

#### Supplementary Fig. 3

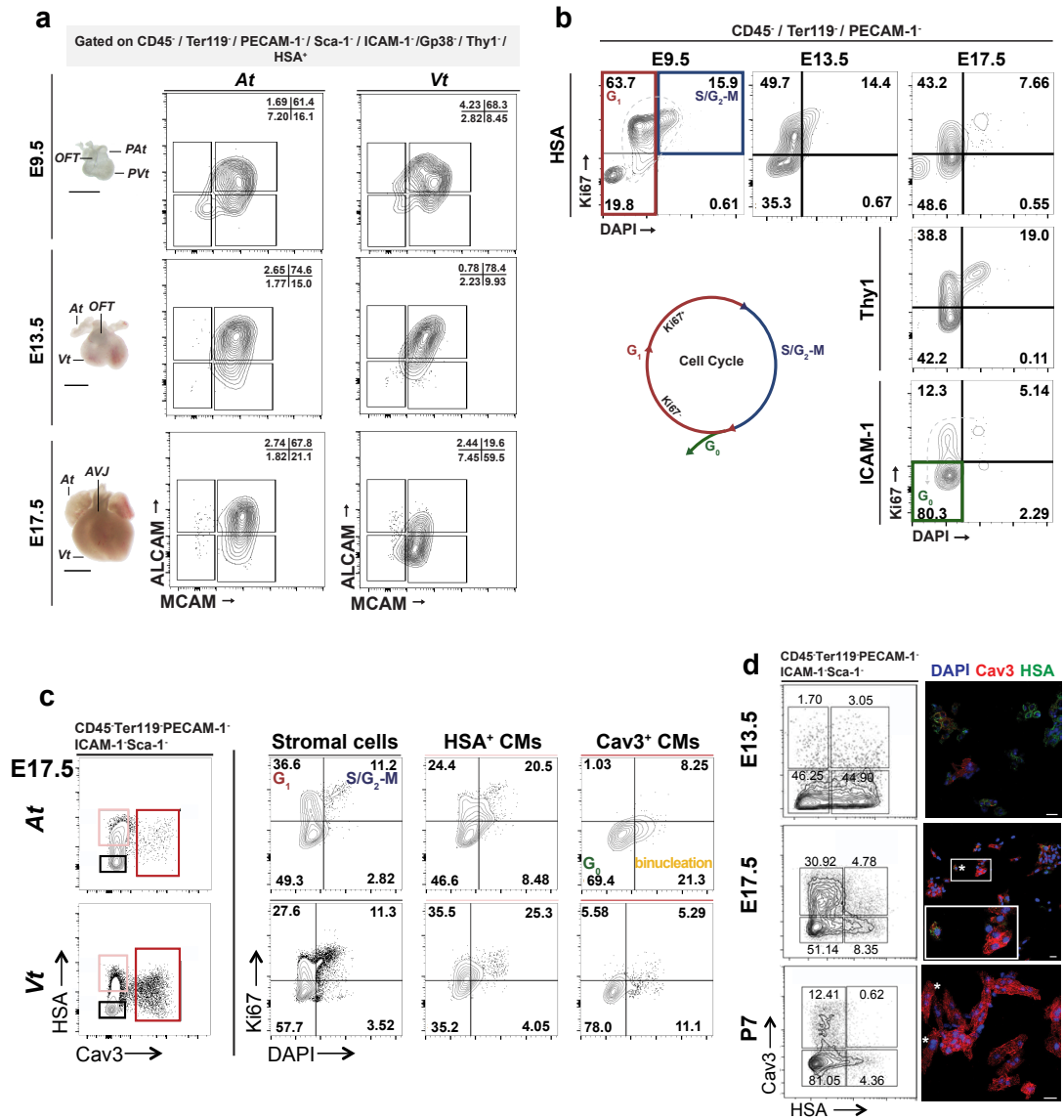

**Supplementary Fig. 3.** Surface phenotype progression of the HSA<sup>+</sup> CMs during heart morphogenesis and their cell cycle status. **a.** Macroscopic view of embryonic hearts at E9.5, E13.5 and E17.5 along with the respective dot plots of flow cytometry data from each heart regions (At or PAAt and Vt or PVt). Scale bar: 1 mm. **b.** Cell cycle analysis of the main cardiac populations combining the surface markers herein identified. Intracellular Ki67 and DAPI allowed determining the frequency of cells in G1 (Ki67<sup>+</sup> and DAPI<sup>2N</sup>, top/bottom left quadrants, red), in S/G2-M (Ki67<sup>+</sup> and DAPI<sup>2N>4N</sup>, top right quadrant, blue) and in G0 (Ki67<sup>-</sup> and DAPI<sup>2N</sup>, bottom left quadrant, green). Contour plots display E9.5 whole heart cells and E13.5 and E17.5 Vt cells. **c.** Cell cycle analysis. G1 (Ki67<sup>+</sup> and DAPI<sup>2N</sup>), S/G2-M (Ki67<sup>+</sup> and DAPI<sup>2N>4N</sup>), G0 (Ki67<sup>-</sup> and DAPI<sup>2N</sup>) and binucleated cells (Ki67<sup>-</sup> and DAPI<sup>4N</sup>) of stromal (black gate), HSA<sup>+</sup>CM (salmon gate) and Cav3<sup>+</sup>CM (red gate) cardiac cells. **d.** HSA and Cav3 expression in E13.5, E17.5 and P7 cardiac cells. Flow cytometry (left panels, n=2) and cytospin (right panels, n=3, 300 cells analyzed in each). Scale bar: 20 μm. At | Atria; Vt | Ventricles.

#### Supplementary Fig. 4

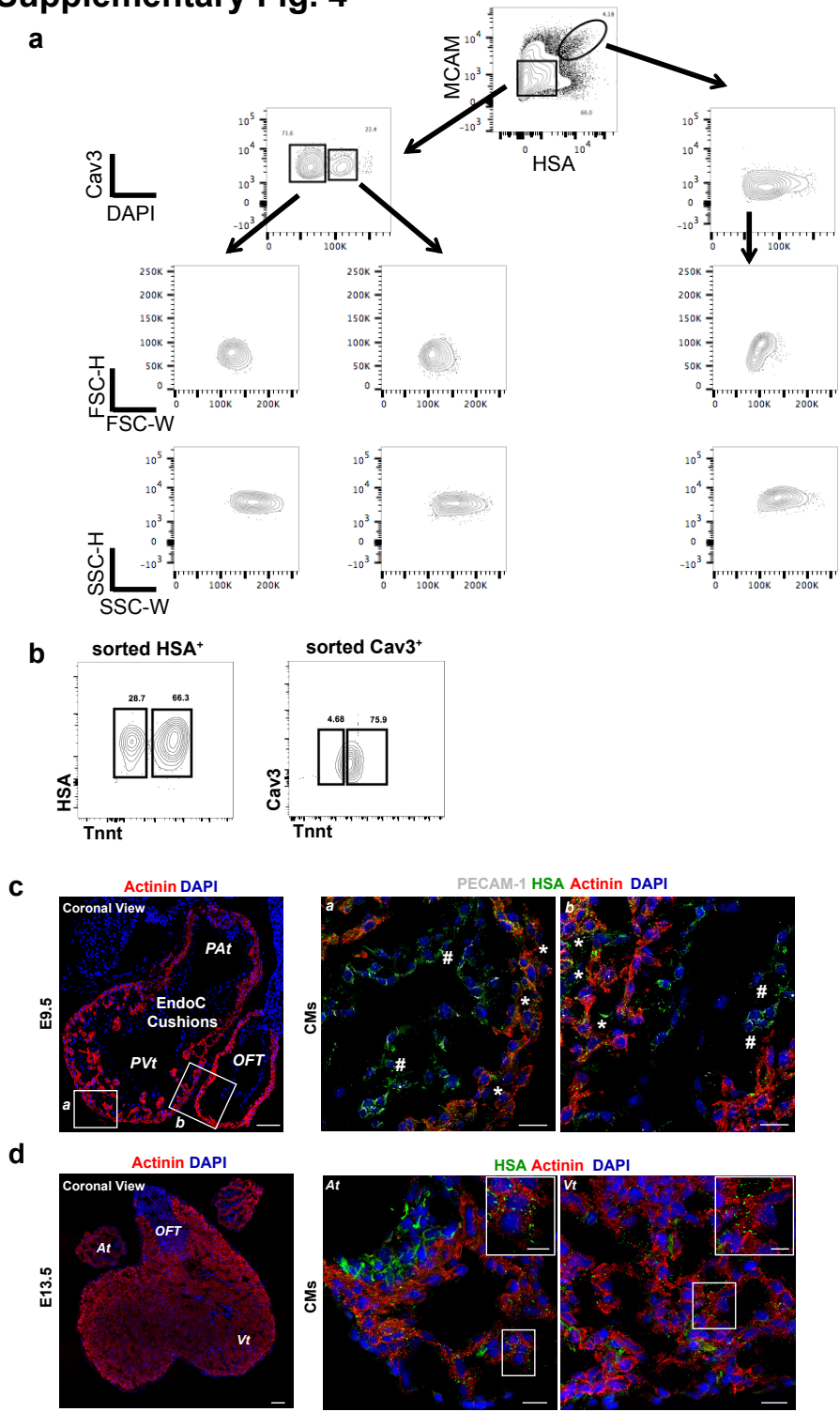

**Supplementary Fig. 4.** Detailed analysis of the two subsets of cardiomyocytes for doublet discrimination and Tnnt expression. **a.** Representative contour plots of the height versus width in the Forward and Side Scatters, excluding the possibility of the 4N subset (binucleated Cav3<sup>+</sup>) to be result of cell doublets. **b.** Demonstration of the Tnnt expression in both HSA<sup>+</sup> and Cav3<sup>+</sup> CM subsets. Because of a technical incompatibility to combine in the same staining Cav3 and Tnnt, we confirmed the presence of Tnnt in the two CM populations (HSA<sup>+</sup> and Cav3<sup>+</sup>) after sorting. **c.** Coronal view of E9.5 heart section stained for Actinin (red) and nuclear content (DAPI; blue), showing the three heart regions. Scale bar: 50 μm. Representative images of E9.5 cardiac tissue display HSA co-expression with either Actinin (CMs, white \*) or PECAM-1 (EndoCs, white #) in the primitive

chambers and EndoC cushions, respectively. Scale bar: 20 $\mu\text{m}$ D. Coronal view of E13.5 heart section stained for Actinin (red) and nuclear content (DAPI; blue), showing the three heart regions. Scale bar: 50 $\mu\text{m}$. Representative images showing CMs ($\text{HSA}^+\text{Actinin}^+$, insets). Scale bars: 20 $\mu\text{m}$ for representative sections and 10 $\mu\text{m}$ for insets. *PAt* | Primitive Atria; *PVt* | Primitive Ventricles; *OFT* | Outflow Tract; EndoC cushions | Endocardial cushions; *At* | Atria; *Vt* | Ventricles.

Supplementary Fig. 5

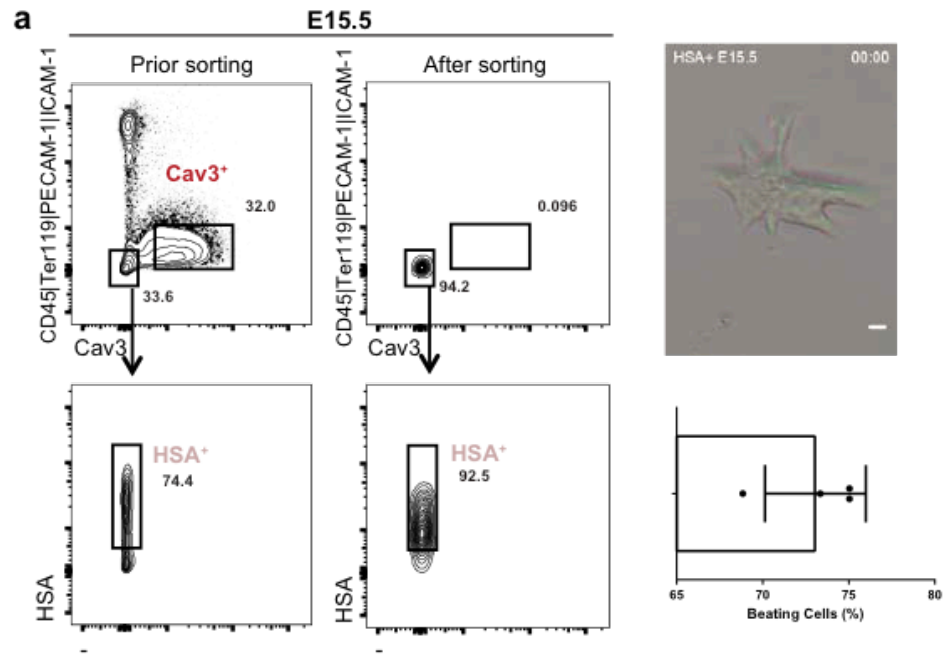

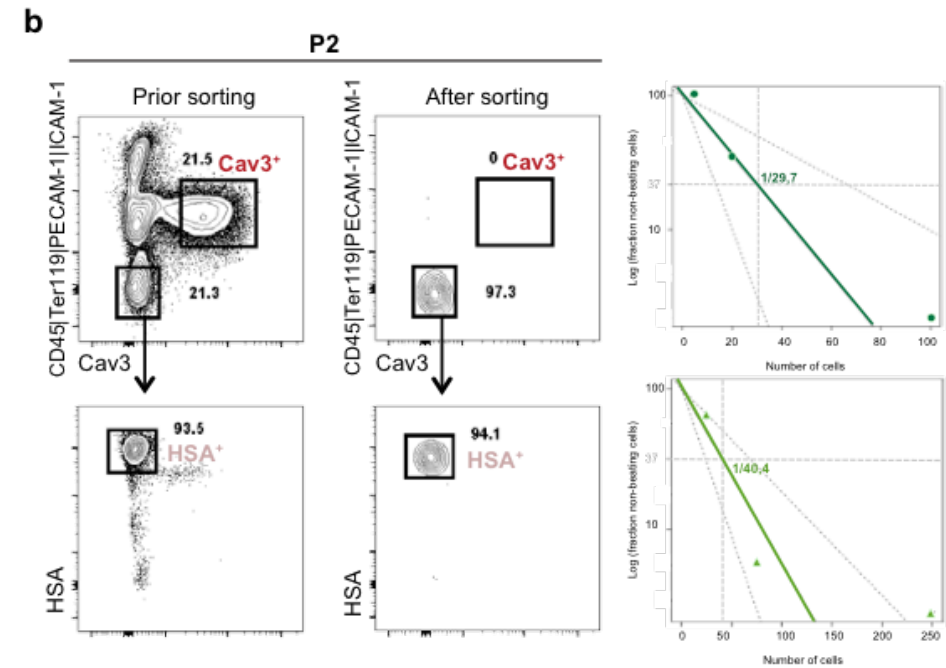

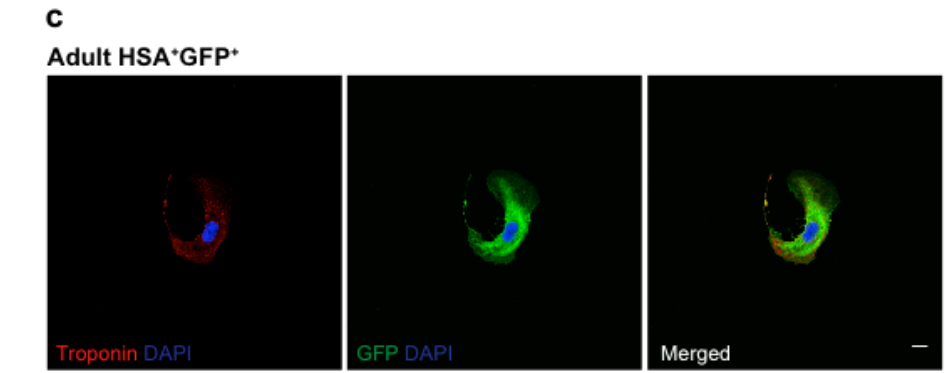

**Supplementary Fig. 5.** Purity analysis and quantification of the HSA<sup>+</sup> CM cultures. **a.** Representative contour plots with the gating strategy to isolate E15.5 HSA<sup>+</sup>Cav3<sup>-</sup> CMs (left plots) and control purity after sorting (right plots). Representative image of a CM in culture (Upper right panel, see also mS1, mS2). Scale: 20μm. Frequency of contractile CMs in cultures (lower right panel). **b.** Representative plots as in **a.** for neonatal and adult cardiac cells (dot plots). Frequency of sorted cells that adhered to gelatin-coated plates (right panels), n=4. Virtually all adherent cells were contractile and expressed cardiac troponin. **c.** Representative image of adult HSA<sup>+</sup> CMs isolated from Ub-GFP mice after 48 hours in culture in 3% O<sub>2</sub>, stained for cardiac troponin (red), GFP (green) and DAPI.

#### Supplementary Fig. 6

**a**

##### Ear-pinna implant experiment design

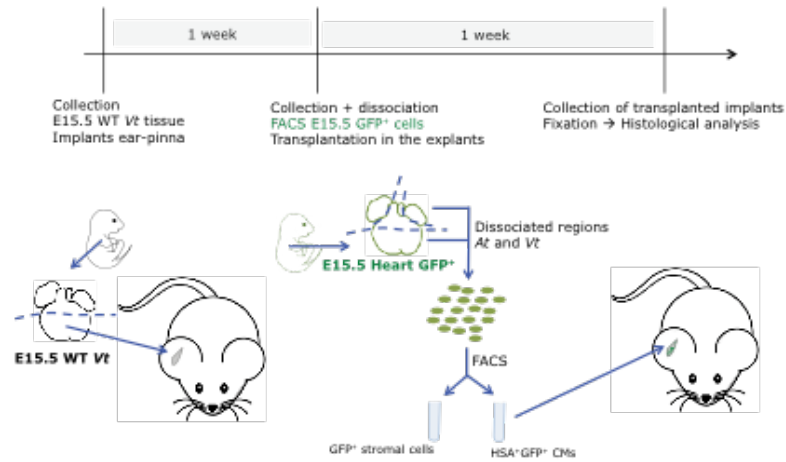

**b**

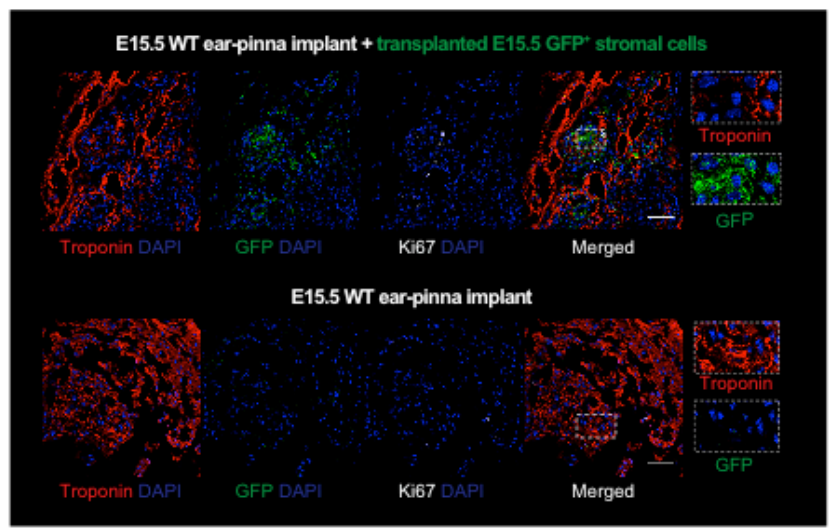

**Supplementary Fig. 6.** Ear-pinna transplantation experiments. **a.** Schematic representation of the experimental design followed in the ear-pinna experiments. **b.** Immunohistochemistry analysis for the expression of Troponin (red), Ki67 (white), DAPI (blue) and GFP (Ub-GFP colonizing cells) of embryonic cardiac tissue implants injected with cardiac stromal cells from the Ub-GFP mice (Upper panels) or implants not injected as controls for the experiment described in Fig 4g. Higher magnification of the region delimited by the white rectangle (right panels). Scale: 50µm.

### Supplementary Fig. 7

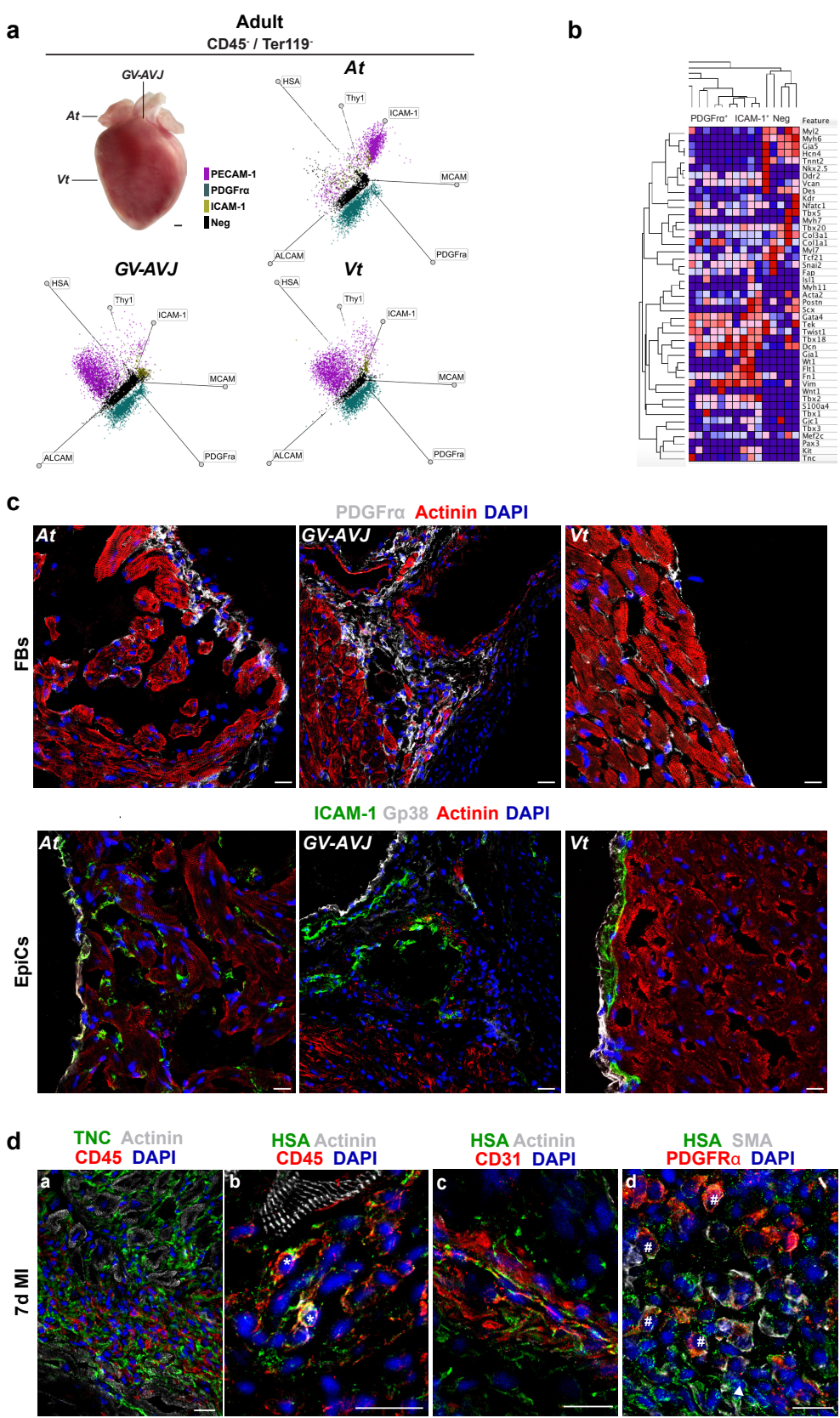

**Supplementary Fig. 7.** Adult cardiac stromal populations. **a.** Macroscopic view of adult heart, depicting the dissected cardiac regions: At, GV-AVJ and Vt. Scale bar: 1 mm. Radar plots of flow cytometry analysis in the CD45<sup>-</sup> and Ter119<sup>-</sup> cell fraction for the surface expression of HSA, Thy1, PECAM-1,

ICAM-1, MCAM, ALCAM and PDGFrα in the indicated heart regions (n=2). **b.** Heat Map displays the unsupervised hierarchical clustering analysis of the multiplex qRT-PCR data at the population level of the indicated adult cardiac populations (100 sorted cells, n=3). Colour-code as in Supplementary Fig. 1. **c.** Representative adult heart sections of the three heart regions (*At*, *GV-AVJ* and *Vt*) stained for Actinin (red) and nuclear content (DAPI; blue), showing 1) interstitial FBs (PDGFrα<sup>+</sup> cells; top row), 2) EpiCs (Gp38<sup>+</sup> cells) and 3) EPDCs (ICAM-1<sup>+</sup> cells); bottom rows. Scale bar: 20 μm. **d.** Representative adult heart sections 7 days post MI stained for Actinin (red) and nuclear content (DAPI; blue), showing a) the cellular infiltrate in the peri-infarcted region (CD45<sup>+</sup> hematopoietic cells and the extracellular matrix protein TenascinC (TNC)); and co-expression of HSA with b) hematopoietic cells (CD45<sup>+</sup>), c) ECs (CD31<sup>+</sup>), and d) SMCs (SMA<sup>+</sup>PDGFrα<sup>+</sup>); Scale bar: 20 μm.

Supplementary Movie S1. E15.5 HSA<sup>+</sup> CM dividing.

Representative live-cell imaging of HSA<sup>+</sup> CM isolated from E15.5 hearts dividing in culture. Time, hr:min. Scale bar: 20μm.

Supplementary Movie S2. E15.5 HSA<sup>+</sup> CM Beating.

Representative example of a contractile HSA<sup>+</sup> CM isolated from E15.5 hearts. Time, min:sec. Scale bar: 20μm.

Supplementary Movie S3. P2 HSA<sup>+</sup> CM Beating.

Representative example of a contractile HSA<sup>+</sup> CM isolated from P2 hearts. Time, min:sec. Scale bar: 20μm.

Supplementary Movie S4. P4 HSA<sup>+</sup> CM Beating.

Representative example of a contractile HSA<sup>+</sup> CM isolated from P4 hearts. Time, min:sec. Scale bar: 20μm.
